# *Plasmodium* rootlet SFA1 coordinates cell replication, apical polarity and cytoskeletal organisation during sexual development

**DOI:** 10.64898/2026.10.30.761637

**Authors:** Declan Brady, Akancha Mishra, Philip Ilani, David J. P. Ferguson, Prince Ubiaru, Robert Markus, Andrew Bottrill, Mohammad Zeeshan, Sue Vaughan, David S. Guttery, Eelco C. Tromer, Anthony A. Holder, Rita Tewari

**Affiliations:** School of Life Sciences, University of Nottingham, Nottingham, UK; Oxford Brookes University, Department of Biological and Medical Sciences, Oxford, UK; School of Life Sciences, Gibbet Hill Campus, University of Warwick, Coventry, UK; Hubrecht Institute-KNAW, Utrecht, The Netherlands; Malaria Parasitology Laboratory, The Francis Crick Institute, London, UK

**Author notes:** Corresponding author (RT). Contributed equally to the work (Declan Brady, Akancha Mishra, Philip Ilani).

**Keywords:** rootlet, striated fibre, SFA1, *Plasmodium*, ookinete, cytoskeleton, apical polarity

## Abstract

Rootlet fibres are evolutionarily conserved cytoskeletal structures involved in basal body positioning and cytoskeletal organisation in diverse eukaryotes. *Plasmodium spps,* undergo complex developmental transitions that require precise coordination of cell polarity, cytoskeletal remodelling and nuclear division. In *Plasmodium* striated fibre assemblin1 (SFA1) and SFA2 have been implicated in linking the centriolar plaque to nascent apical poles during schizogony and sporogony. However, their role in sexual development is unclear. Here we show SFA1 is absent from the male basal body but is present in female gametocytes, where it marks the polarity initiation site during zygote-to-ookinete differentiation. We also identify associations between SFA1, SFA2 and pellicle-associated proteins. Loss of *sfa1* disrupts apical polarity, pellicle organisation and subpellicular microtubules, producing abnormal immotile ookinetes and affecting further sporozoite development, leading to complete loss of parasite transmission. Together, these findings support an important role for SFA1 in organising the cytoskeletal architecture required for malaria transmission.

## Introduction

Rootlets are evolutionarily conserved, cytoskeletal fibres that contribute to many cellular functions including ciliary and flagellar motility, basal body positioning, centrosome cohesion, centriole positioning, and mechanosensory signalling (*1*). Rootlets are closely associated with basal bodies in flagellated organisms, including *Chlamydomonas*, where they are important in cellular architecture and symmetry (*2, 3*). Loss of rootlets has been associated with mitotic defects, as centrosome disjunction during cell division is closely linked to rootlet splitting and reorganisation (*1, 4, 5*). In many unicellular eukaryotes, particularly species lacking cilia, rootlets are formed from striated fibre assemblins (SFAs) (*6*) which have been found in a wide range of organisms, including the phylum *Apicomplexa* (*7*).

Apicomplexa are a diverse group of obligate intracellular parasites, including *Plasmodium* spp., the causative agents of malaria, and *Toxoplasma gondii*, the causative agent of toxoplasmosis. The invasive stages of these parasites are highly polarised, with an apical complex located at the anterior end of the cell, and this specialised structure is essential for gliding motility, host-cell recognition, and invasion (*8*). Motility is not driven by cilia or flagella but by the glideosome, an actomyosin-based motor complex. The glideosome is anchored within the pellicle, comprised of the plasma membrane and a flattened vacuolar structure called the inner membrane complex (IMC) (*8, 9*). The pronounced apical polarity requires precise cytoskeletal organisation that is also linked to the machinery of cell division and proliferation.

*Plasmodium* has a complex life cycle, alternating between vertebrate and mosquito hosts and including three morphologically distinct invasive zoite stages: merozoite, ookinete, and sporozoite. These stages invade host tissue and cells and subsequently develop into proliferative forms characterised by atypical modes of mitosis and meiosis during cell division (*10*). Only one stage of the *Plasmodium* life cycle, male gametogenesis, has basal bodies and formation of flagella (*10, 11*).

Three SFA proteins have been identified in *T. gondii* and shown to be essential for establishing cell polarity through association with both the conoid in the apical complex and the cell division machinery (*12*). In *Plasmodium*, two homologues, SFA1 and SFA2, have been described as present and functioning during two proliferative stages: schizogony that gives rise to merozoites within mammalian erythrocytes, and sporogony that produces sporozoites, in oocysts within the mosquito vector (*13*). These developmental programmes are both characterised by extensive asexual mitosis and cell replication. The sexual stage of the life cycle is initiated by activation of gametocytes in a mosquito blood meal and formation of gametes. Following fertilisation, the diploid zygote differentiates into a polarised and motile ookinete within the mosquito midgut. The ookinete is a unique invasive stage during the life cycle as it develops post-fertilisation from the round zygote, differentiating and elongating extracellularly within the midgut of the mosquito vector, invading the gut wall and colonising intercellular space to produce an oocyst (*10, 14-17*). It is also different from other stages in that it does not arise due to multiplication within a coenocyte such as a schizont or oocyst and does not form a parasitophorous vacuole following invasion of the gut wall. During this developmental transition one round of DNA replication and meiosis occur, prior to the ookinete migrating through the mosquito gut wall and forming an oocyst (*10, 17*). To date, the localisation and function of SFAs during ookinete development have not been fully characterised.

Here, we investigate the spatiotemporal location and function of *Plasmodium* SFA1 (PBANKA_1227000) during parasite development across multiple stages of the *Plasmodium* life cycle, with greater focus on ookinete differentiation and development using live-cell imaging, expansion microscopy, reverse genetics and protein interaction studies. Our results reveal an unexpected and essential role for SFA1 during zygote-to-ookinete differentiation and its absence in basal body or flagella formation during male gametogenesis. These findings demonstrate that malaria parasites have repurposed a conserved cytoskeletal component of rootlet to support developmental progression and parasite transmission through the mosquito vector.

## Results

### SFA1 is present in developing invasive and proliferative stages of the parasite life cycle but absent during male gametogenesis

To investigate the spatiotemporal location of SFA1 throughout the *P. berghei* life cycle, the endogenous *sfa1* locus was modified to tag the C-terminus of SFA1 with green fluorescent protein (GFP). Correct integration of the *gfp* sequence and expression of the fusion protein were confirmed by diagnostic PCR and western blot analysis, respectively (Supplementary Figure 1A-C). Imaging of the SFA1-GFP fusion protein in live cells was performed for multiple parasite stages in both mammalian and mosquito hosts, but with a specific emphasis on the zygote to ookinete transition.

During asexual blood-stage parasite development, SFA1-GFP was not detected in ring or trophozoite stages (Supplementary Figure 1D), but following the onset of nuclear division and schizogony, SFA1-GFP fluorescence was visible as a discrete punctum adjacent to individual nuclei. During the asynchronous nuclear division, prior to division of each nucleus SFA1-GFP appeared as duplicate foci. This SFA1-GFP duplication preceded successive rounds of nuclear division throughout schizogony, but at the end of the process SFA1-GFP fluorescence was absent from mature segmented schizonts containing differentiated merozoites (Figure 1A, Supplementary Figure 1D).

**Figure 1.**
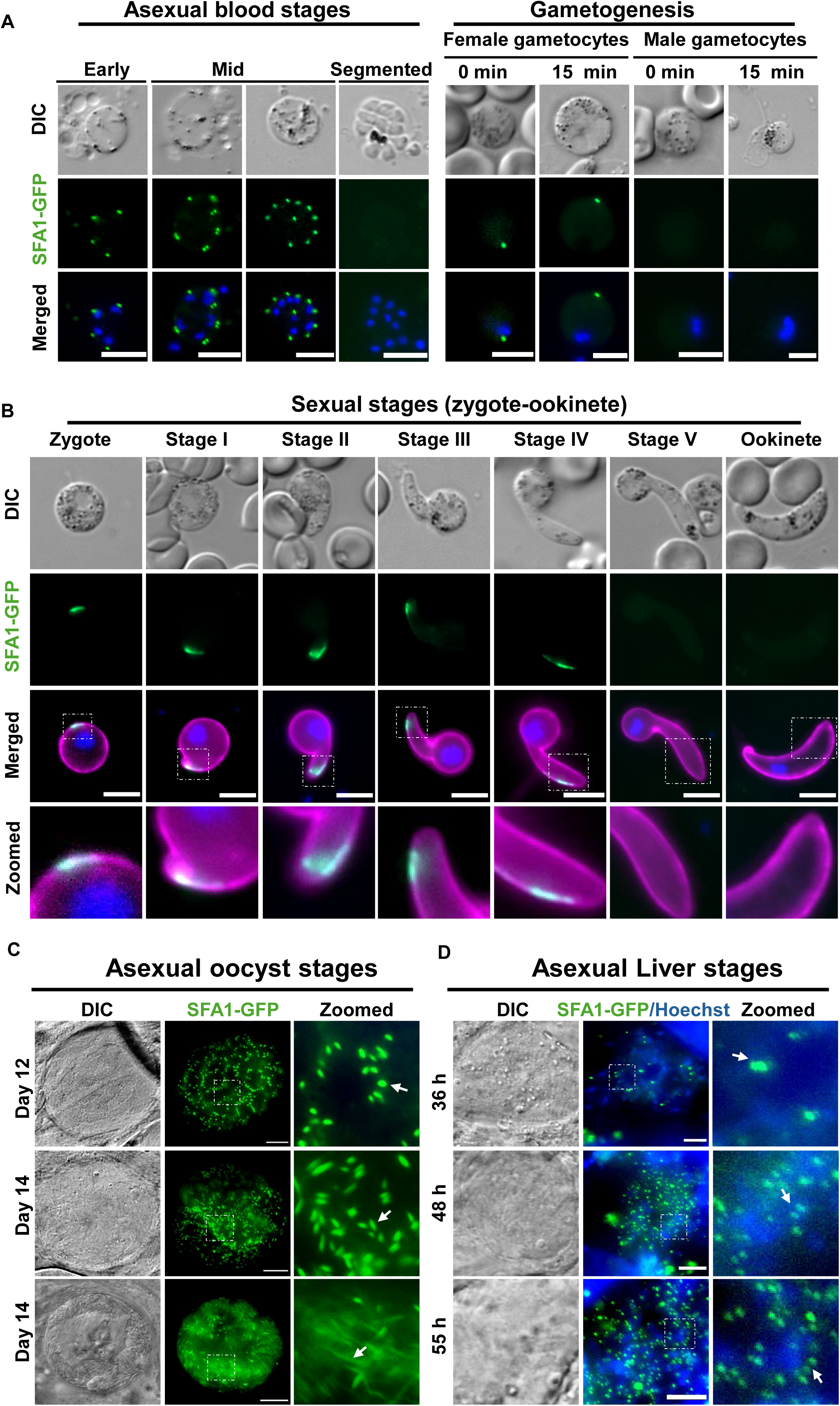
Spatiotemporal location of SFA1 in *Plasmodium berghei* life cycle: **(A)** SFA1-GFP (green) expression in asexual blood stages from early on the left to mid to segmented schizont. A female and a male gametocyte are shown on the right side of the figure. The top row of panels are differential interference contrast (DIC), the middle row are GFP (green) fluorescence, and the bottom row are merged GFP fluorescence and Hoechst stain (blue) for DNA. Scale bar 5 µm. **(B)** Post-fertilisation expression pattern of SFA1-GFP (green) from the zygote to the mature ookinete. The top row of images are DIC, the second row of images are GFP fluorescence, the third row of images are merged GFP, Hoechst blue (DNA), P28 surface marker, with regions of the image magnified in the lower panels indicated by the white boxes. Insets show zoom of apical localisation. Scale bar 5 µm. **(C)** SFA1-GFP expression in the oocyst (asexual mosquito stages) (green), with dot-like appearance in the early sporoblast stage, and elongated expression in the sporozoite segregation stages. Left side panels are DIC, other panels are GFP). Insets show zoom of the SFA1-GFP expression pattern in oocyst stages. Scale bar 10 µm. Arrow indicates the SFA1 (green) localisation **(D)** SFA1-GFP (green) expression in liver stages at 36, 48 and 55 hpi. Hoechst stain (blue) for DNA. Insets show zoom of SFA1-GFP expression pattern in oocyst stages. Scale bar 5 µm. The inset shows the magnified view of the boxed region. Arrow indicates the SFA1 (green) localisation.

During sexual development, we observed distinct foci of SFA1-GFP in non-activated female gametocytes, whereas no signal was detected in non-activated male gametocytes (Figure 1A, Supplementary figure 1D). Following gametocyte activation, SFA1-GFP fluorescence remained in female gametocytes at the cell periphery and was absent from activated male gametocytes despite the formation of basal bodies and flagella at this stage (Figure 1A). Following fertilization, the SFA1-GFP signal persisted at the cell periphery in the early zygote but during morphological transformation and elongation into an ookinete, it was concentrated at the developing apical end of the cell. In the early ookinete developmental stages (I–III) (*18*) the protein had an elongated location along one side of the nascent apical protrusion (Figure 1B, Supplementary Figure 1E). During later stages of ookinete development, the fluorescence signal started diminishing in stage IV ookinetes and was completely absent from stage V and mature ookinetes (Figure 1B, Supplementary Figure 1E), reminiscent of the absence of the protein from mature schizonts. These data indicate that the location of SFA1 is restricted to processes in the differentiation of ookinetes rather than at the fully differentiated ookinete invasive stage.

To investigate the location of SFA1-GFP during sporogony, mosquitoes were infected with SFA1-GFP parasites and the development of oocysts and sporozoites was monitored over 21 days. No SFA1-GFP fluorescence was detected in early oocysts at day 7 post-infection (Supplementary Figure 1F). However, during sporoblast development at between days 12 and 14, SFA1-GFP was prominent as discrete foci that subsequently extended into elongated structures as sporulation progressed (Figure 1C). By day 21 post-infection, most oocysts had reached advanced stages of development, and SFA1-GFP fluorescence was absent from the majority of oocysts (Supplementary Figure 1F). In sporozoites isolated from developing oocysts most had no detectable SFA1-GFP fluorescence, although a small proportion (< 1%) retained a signal (Supplementary Figure 1G, H). Similarly, no SFA1-GFP fluorescence was observed in salivary gland sporozoites indicating that SFA1 is lost as the sporozoite matures (Supplementary figure 1G,H).

Finally, the presence of SFA1-GFP was examined during liver-stage development following hepatocyte infection with sporozoites. No SFA1-GFP was detected at approximately 12-24hours post-infection (Supplementary Figure 1I), during early liver-stage development, but in the subsequent schizont stage undergoing active mitotic nuclear division at 36-55 hpi, a prominent signal was detected (Figure 1D). Consistent with the asynchronous pattern of nuclear division, SFA1-GFP initially appeared as single nucleus-associated foci, whereas paired foci were observed at later stages of extensive nuclear multiplication. This pattern suggests that SFA1-GFP focus duplication accompanies nuclear division, as observed during asexual blood-stage schizogony. Similarly, the SFA1-GFP fluorescence disappeared during the terminal differentiation of exoerythrocytic merozoites (Supplementary Figure 1I).

These observations demonstrate that SFA1 has a dynamic, developmentally regulated subcellular location throughout the *P. berghei* life cycle. Its expression is predominantly associated with proliferative stages during nuclear division, followed by its disappearance upon terminal differentiation of the cell. The SFA1 signal is duplicated prior to each nuclear division during schizogony, suggesting an association with the machinery coordinating nuclear division and cell-cycle progression. Interestingly, it is not detected during male gamete formation, which requires three rapid rounds of mitosis and synthesis of basal bodies and flagella.

### SFA1 associates with the apical subpellicular microtubule organising centre (MTOC) during nuclear division and ookinete development

Based on the location of SFA1-GFP in proliferative parasite stages and its known association with the centrosome or spindle pole body in other eukaryotes, we investigated the spatial relationship of SFA1-GFP to the microtubule-organising and chromosome-segregation machineries. We used co-localisation studies with antibodies to centrin and α-tubulin, and generated parasites expressing both SFA1-GFP and the kinetochore marker NDC80-mCherry.

Asexual blood-stage schizonts expressing SFA1-GFP were co-stained with anti-centrin and anti-α-tubulin antibodies, revealing that SFA1-GFP is located adjacent to the outer centriolar plaque (CP)/spindle microtubule-organizing centre (MTOC) (Figure 2A). Structured illumination microscopy (SIM) revealed that SFA1-GFP and CP duplication are coincident. Following SFA1-GFP duplication, nuclear division occurred, with each daughter nucleus retaining one CP and one SFA1-GFP signal (Figure 2A). This pattern was observed throughout schizogony, as different nuclei within the same cell underwent asynchronous successive rounds of independent division. Staining the SFA1-GFP cells with α-tubulin antibodies showed that the mitotic spindle originates from the MTOC immediately beneath the SFA1-GFP signal, placing SFA1 in close proximity to the spindle pole (Figure 2A).

**Figure 2.**
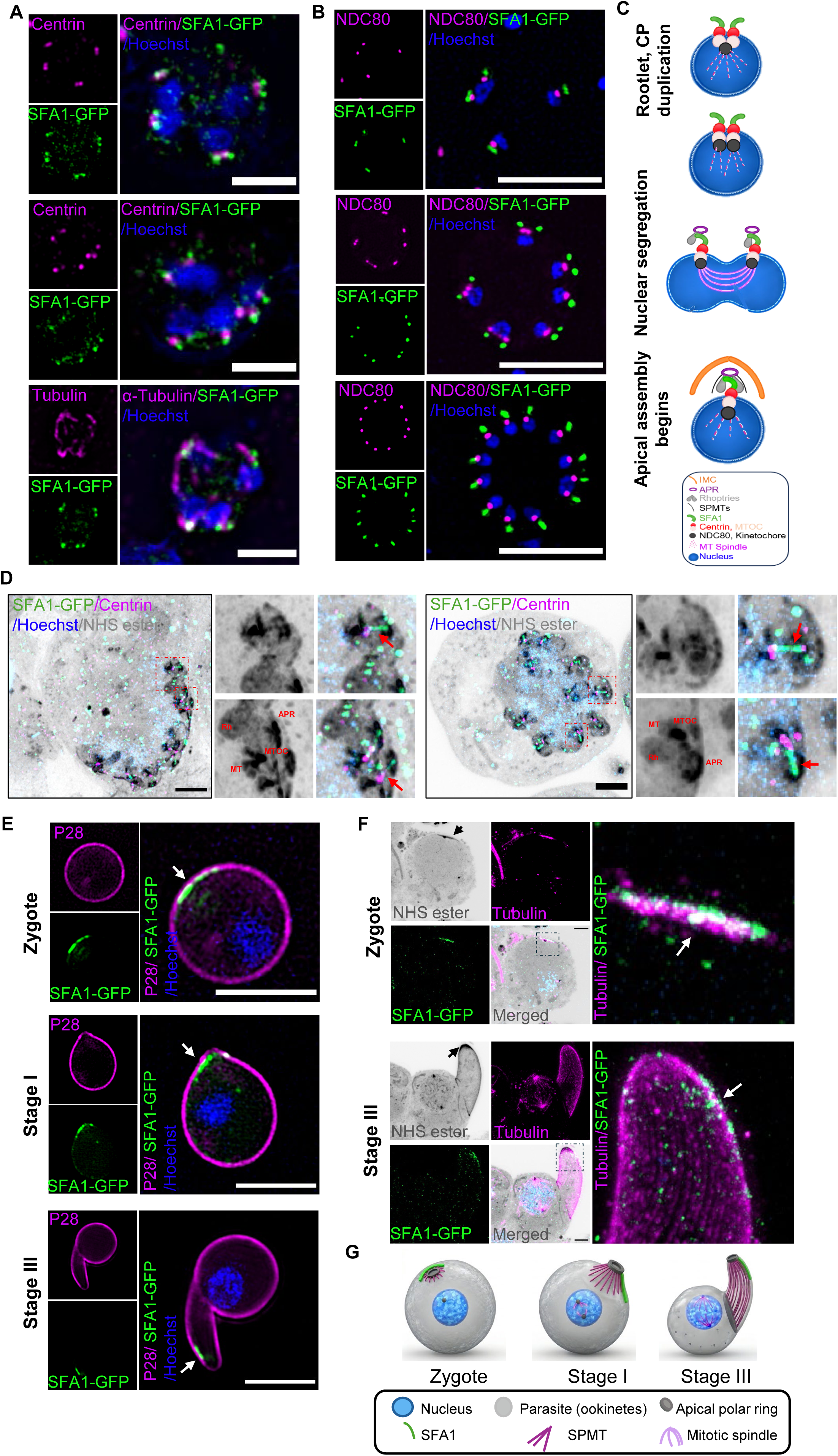
Ultrastructure imaging of SFA1-GFP in asexual and sexual stages: **(A)** Immunofluorescence staining of SFA1-GFP schizonts, co-stained with anti-GFP (green) and anti-Centrin (magenta; first and second panel), and anti-GFP (green) and tubulin (magenta; third panel), Hoechst (blue) imaged by Structured Illumination Microscopy (SIM). Scale bar 5 µm. **(B)** Structured Illumination Microscopy (SIM) imaging of dual tagged parasite line. NDC80 cherry (magenta; kinetochore) and SFA1-GFP (green) stained with Hoechst (blue) showing the asynchronous nuclear division during early schizogony. Scale bar 5 µm. **(C)** Cartoon schematic of early schizogony events showing segregation of SFA1 (green) along with Centrin (red; outer centriolar plaque CP) and NDC80 (black; kinetochore), nucleus (blue) during asynchronous nuclear division. Legends are highlighted below. **(D)** Ultrastructure Expansion Microscopy (U-ExM) imaging of SFA1-GFP schizonts co-stained with anti-GFP (green), anti-Centrin (magenta) antibody, NHS ester dye (grey) and Hoechst (blue). Scale bar 5 µm. The inset shows the magnified view of the boxed region. Red arrow indicates the SFA1 fibre (green). MT represents mitotic spindle, Apical polar region APR, Rh rhoptries, MTOC Microtubule organising centre. **(E)** SIM imaging of live SFA1-GFP (green) ookinetes stages co-stained with P28 antibody (magenta) and Hoechst (blue) at 4-9 h post activation (hpa). Scale bar 5µm. Arrow indicates the SFA1 (green) localization. **(F)** U-ExM imaging of SFA1-GFP ookinetes fixed at 4-9 hpi, co-stained with anti-GFP (green) and anti-tubulin (magenta) antibody, NHS ester dye (grey) and Hoechst (blue), highlighting the expression in the zygote stage ( top panel) and stage III ( bottom panel). Scale bar 5 µm. The inset shows the magnified view of the boxed region. Arrow indicates the SFA1 (green) localization. **(G)** Cartoon schematic of SFA1 spatiotemporal location in the developing ookinete stages.

To further examine the spatiotemporal relationship of SFA1 with the chromosome-segregation machinery, we analysed parasites expressing both SFA1-GFP and the kinetochore marker, NDC80-mCherry. SFA1-GFP was consistently observed in close proximity to the NDC80-labelled kinetochore, but the two signals were distinct and did not overlap, indicating that SFA1 is not a component of the kinetochore. Despite the spatial proximity, the temporal pattern of division was distinct: SFA1-GFP was duplicated first, followed by duplication of NDC80-labelled kinetochores, and subsequent nuclear division (Figure 2B). At different stages of schizogony, asynchronous progression through this sequence occurred until nuclear division was complete, with an individual NDC80-labelled kinetochore signal and a single SFA1-GFP signal associated with each nucleus (Figure 2B and C). However, in mature segmented schizonts, SFA1-GFP fluorescence was undetectable (Supplementary Figure 1D), indicating that SFA1 is primarily associated with actively dividing nuclei during schizogony and is lost upon completion and merozoite formation.

To examine the spatial relationships between SFA1, the apical end of the cell, and the CP, we used ultrastructure expansion microscopy (U-ExM) with anti-GFP and anti-centrin antibodies together with NHS-ester staining to visualise protein-dense cellular structures. This analysis revealed a thin SFA1-GFP fibre extending from the apical region towards the outer CP, suggesting a continuous physical connection (Figure 2D). This architecture was observed both in parasites undergoing nuclear division and segregation (Figure 2D, left panel), as well as in more mature cells where nuclear division and apical organisation were largely complete (Figure 2D, right panel). Importantly, the SFA1-outer CP connection was evident throughout these stages, suggesting that the association is maintained during daughter-cell development prior to cytokinesis (Figure 2D).

During the sexual development stage, we examined the location of SFA1 during zygote to ookinete differentiation using SIM imaging of SFA1-GFP parasites labelled with an antibody to the ookinete surface marker P28 (*19*). In zygotes and during early ookinete development, SFA1-GFP was present as an arc-like structure located toward the developing apical region. In stage I ookinetes, the signal was concentrated at the apical end, whereas by stage II-III the SFA1-GFP signal appeared to form a more elongated fibre-like structure extending from the apical region (Figure 2E). However, by stage IV, the signal had begun to diminish and was not detectable in stage V or mature ookinetes as described earlier (Figure 1B).

High resolution analysis of SFA1 architecture during ookinete differentiation was obtained using U-ExM with parasites fixed at 7-9 h post-activation and stained with anti-GFP, anti-α-tubulin and NHS ester. In zygote and stage I ookinetes, SFA1-GFP was located in close proximity to the apical polar ring and extended along the subpellicular region marked by NHS-ester and microtubule staining (Figure 2F, top panel, Supplementary Figure 2A). In stage II-III ookinetes, SFA1-GFP had an elongated fibre-like morphology from the apical end of the cell, which extended along the developing apical-subpellicular region (Figure 2F, bottom panel, Supplementary Figure 2A). Collectively, our observations place SFA1 in close association with the apical subpellicular MTOC and the apical cytoskeletal architecture during both asexual nuclear division and ookinete morphogenesis (Figure 2G).

### SFA1 forms a complex with SFA2 and other apical proteins

To identify proteins associated with SFA1 during parasite development, we performed GFP immunoprecipitation followed by mass spectrometry (IP–MS) in schizonts, non-activated gametocytes and stage-II/III ookinetes (7-9h post activation) (*18*) (Figure 3). SFA2 (PBANKA_1012000) was consistently recovered in all three stages, supporting its previously established association with SFA1(*13*).

**Figure 3.**
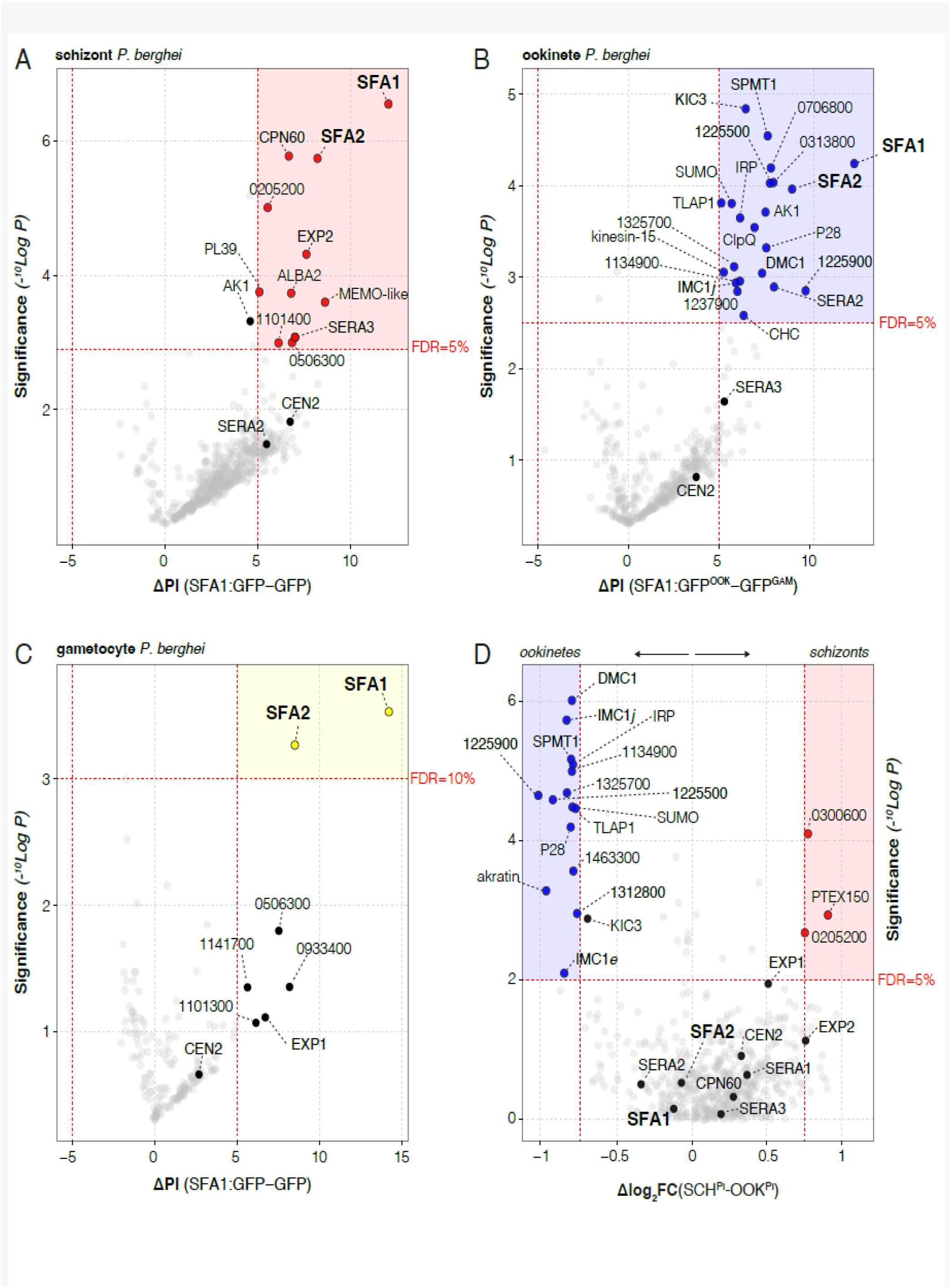
SFA1-GFP immunoprecipitation recovers SFA2 and proteins associated with developing ookinete apical architecture. **(A–C)** Volcano plots comparing crosslinked SFA1-GFP and GFP-only immunoprecipitations from **(A)** schizonts, **(B)** developing ookinetes and **(C)** non-activated gametocytes. Precursor intensities were normalised by the total identified precursor intensity within each run and log₂-transformed. Plotted proteins were detected in at least two retained SFA1-GFP preparations. Positive ΔPI values indicate higher mean log₂-normalised precursor intensity in the SFA1-GFP group. Replicate numbers (SFA1-GFP/control) were 4/4 for schizonts, 3/3 for ookinetes and 2/3 for gametocytes. For panel B, gametocyte GFP-only samples were used as controls, rather than stage-matched ookinete controls (due to lack of signal). The vertical axis shows −log₁₀(P) from the protein-level Welch test. Multiple correction following the Benjamini-Hochberg procedure. FDR set at 5%; 10% for gametocytes. **(D)** Comparison of proteins recovered in SFA1-GFP pulldowns from schizonts and ookinetes, based on normalised precursor intensities. Negative values indicate preferential recovery in ookinetes, whereas positive values indicate preferential recovery in schizonts. Selected proteins associated with the apical cytoskeleton, inner membrane complex (IMC), and other cellular compartments are annotated, including members of the serine repeat antigen (SERA) family. Dashed horizontal lines indicate the FDR thresholds marked in each panel (5% in A, B and D; 10% in C).

Developing-ookinetes pulldowns preferentially recovered (Figure 3B) proteins associated with the apical cytoskeleton or pellicle. Specifically, these included proteins related to subpellicular microtubule protein 1 (SPMT1; PBANKA_0810700): TLAP1 (PBANKA_1444800), IMC1j (PBANKA_1120400), IMC1*e* (PBANKA_0402700), KIC3 (PBANKA_0815400), and PBANKA_1225500 (ortholog in *P. falciparum*: PF3D7_0805100) (*20*) that were reproducibly detected in ookinete SFA1-GFP preparations but not in the corresponding schizont pulldowns (Figure 3D). A conserved protein of unknown function (PBANKA_1225900) showed a similar distribution, being among the most abundant proteins recovered from developing ookinetes. The preferential recovery of these proteins is consistent with the location of SFA1 along the developing apical region and subpellicular microtubule array.

We also identified two members of the serine-repeat antigen family, SERA2 (PBANKA_0305000) and SERA3 (PBANKA_0304900), in both schizont and ookinete pulldowns. SERAs are associated primarily with parasite proteolytic and secretory processes. SERA3 has recently been localised to the ookinete surface and shown to undergo proteolytic processing at this stage (*21*); whereas SERA2 was previously characterised predominantly in late blood and liver stages (*22*). The recovery of both proteins suggests that the SFA1 pulldowns also capture proteins associated with the developing parasite surface or secretory compartments. Whether these represent specific SFA1-associated proteins or proteins recovered through their proximity to the developing pellicle remains to be established.

### SFA1 is essential for ookinete development, cytoskeleton organisation and parasite transmission through the mosquito

To study the function of SFA1 throughout the parasite life cycle, we made a *sfa1* knockout parasite line (*sfa1*-KO) using conventional gene deletion approaches (*23*) and examined the resultant phenotype. Successful gene deletion was confirmed using integration PCR at the *sfa1* locus and by qPCR of asexual blood-stage parasites (Supplementary Figure 3A-C).

First, we assessed the effect of *sfa1* deletion on asexual blood-stage parasite growth. Mice were infected intraperitoneally with 1,000 parasites and parasitaemia was monitored daily by microscopy with Giemsa-stained blood smears. Wild-type (WT) parasites established a patent infection by day 4, whereas with *sfa1*-KO parasites there was a modest but reproducible delay (Figure 4A). This may be due to a replication defect in some cells.. However, in contrast to previous studies (*13*), both control and mutant produced similar numbers of merozoite-containing nuclei. Nevertheless, infected mice eventually developed severe disease and were euthanised.

**Figure 4.**
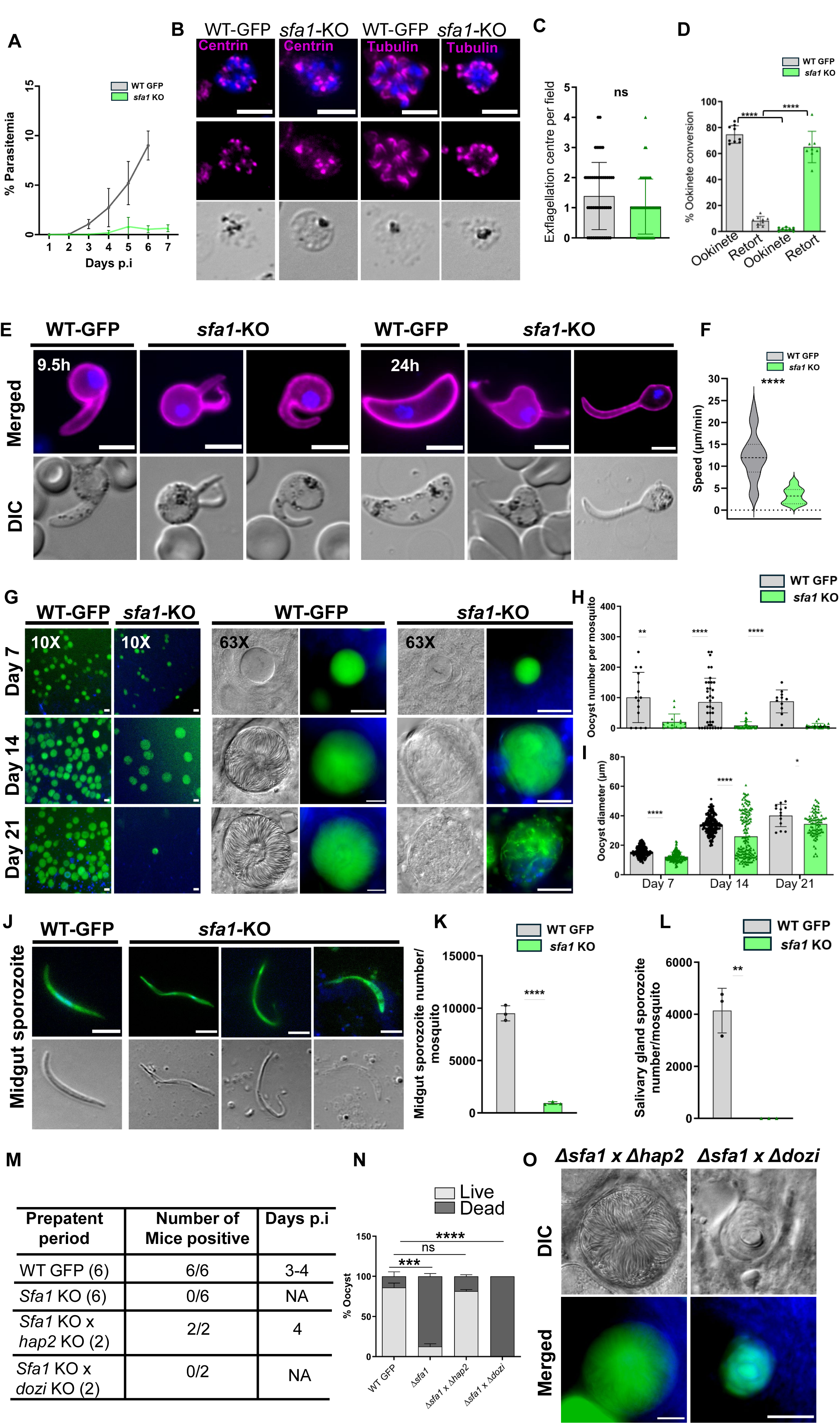
Phenotypic analysis of *sfa1*-KO parasites in *P. berghei* life cycle: **(A)** Percentage parasitaemia growth rate of WT-GFP and *sfa1**-***KO parasites. The plot shows the mean ± SD, unpaired t-test, three independent experiments. **(B)** Immunofluorescence imaging of WT-GFP and *sfa1* KO blood stage schizonts co-stained with centrin (magenta), tubulin (magenta) and Hoechst (blue). Scale bar 5 µm. **(C)** Exflagellation centres per field at 15 mins post-activation. n=3 independent experiments (> 7 fields per experiment). ns, not significant, *P*=0.0950, two tailed unpaired t-test. **(D)** Percent ookinete and retort conversion of WT-GFP and *sfa1**-***KO parasites at 24 h post activation (hpa). Mean ± SD, two-tailed unpaired t-test, ****p<0.0001, ****p<0.0001, n=4 independent experiments. **(E)** Live cell imaging of WT-GFP and *sfa1**-***KO ookinetes at 9.5 and 24 hpa. Parasites stained with P28 antibody (magenta) and Hoechst (blue). Scale bar: 5 µm. **(F)** Motility assay of WT and *sfa1* KO ookinetes at 24 hpa, indicating speed (µm/min). Mean ± SD, two-tailed Mann-Whitney test, ****p<0.0001, n=2 independent experiments. **(G)** Live cell imaging of midguts infected with WT-GFP or *sfa1**-***KO parasites at 7-, 14- and 21-days post-infection at 10X and 63X magnification. (n>20, 3 independent experiments). Scale bar: 10 µm. **(H)** Total GFP-positive oocysts in *sfa1**-***KO-infected midguts compared to WT-GFP at 7-,14-, and 21 dpi. Mean ± SEM, two-tailed unpaired t-test, **p=0.0012, ****p<0.0001. n=5 independent experiments. **(I)** Diameter of GFP-positive oocysts at 7-, 14-, and 21 dpi. Mean ± SEM, two-tailed unpaired t-test, ****p<0.0001,*p=0.0174. n=5 independent experiments. **(J)** Live cell imaging of WT GFP and *sfa1* KO sporozoites at 14 dpi. Sporozoites were stained with Hoechst (blue). (n>20, three independent experiments). **(K)** Midgut sporozoite number of *sfa1**-***KO-infected guts compared to WT-GFP infected guts at 14 dpi. Mean ± SEM, two-tailed unpaired t-test, ****p<0.0001. n=3 independent experiments. **(L)** Salivary gland sporozoite number obtained from WT GFP and *sfa1* KO infected mosquitoes. Mean ± SEM, two-tailed unpaired t-test, p=0.0011, n=3 independent experiments. **(M)** Pre-patent period observation of *sfa1-*KO compared with WT-GFP parasites (n=6 independent experiments) and *sfa1* KO x *hap2* KO and *sfa1* KO x *dozi* KO parasites in infected mice (n=2 independent experiment). **(N)** Percentage of oocyst (sporulation) observed in WT GFP and *sfa1* KO infected midguts and percentage of oocyst (sporulation) rescued with cross of *sfa1* KO x *hap2* KO and *sfa1* KO x *dozi* KO infected midguts. P=0.0002, ns p=0.5798, ****p<0.0001, two-way ANOVA, two independent experiments. **(O)** Live cell imaging of midguts infected with *sfa1* KO x *hap2* KO and *sfa1* KO x *dozi* KO parasites at day 14-post-infection at 63X magnification. (n>20, 2 independent experiments). Scale bar: 10 µm.

To investigate potential cellular defects underlying the growth delay, we performed immunofluorescence analysis of schizonts co-labelled with anti-centrin and anti-tubulin antibodies. In early dividing WT-GFP schizonts, centrin foci were closely associated with individual nuclei, and tubulin staining revealed discrete microtubule structures associated with dividing nuclei. By contrast, a subset of *sfa1*-KO schizonts exhibited an altered distribution of centrin foci relative to the nuclei: some nuclei lacked an associated detectable centrin focus, whereas others were associated with multiple foci (Figure 4B, Supplementary Figure 3D, E). In addition, tubulin staining appeared more focused in mutant schizonts than in WT controls (Figure 4B, Supplementary Figure 3D, F). Although the distribution of centrin-associated structures and tubulin staining was altered in a subset of *sfa1*-KO schizonts, the numbers of merozoite-containing nuclei were similar to those in the WT parasites. Together, these observations suggest that *sfa1* deletion perturbs the spatial organisation of centrin-associated structures and microtubules during various stages of schizogony.

During sexual development, WT and *sfa1*-KO parasites produced comparable numbers of gametocytes (4-6%) and following male gametocyte activation and gametogenesis no significant difference in exflagellation was observed between the two lines (Figure 4C). However, following fertilisation and development from zygote to ookinete, WT parasites differentiated into mature ookinetes, at 24 h post-activation, whereas approximately 90% of *sfa1*-KO parasites failed to completely differentiate. Instead, they formed abnormal retort-stage intermediates, resulting in a significant reduction in ookinete conversion (*P* < 0.001, Figure 4D).

To determine when this developmental defect arose, we examined ookinete development using live cell imaging. At approximately 9.5 h post-activation WT parasites were at approximately stage II-III, whereas *sfa1*-KO parasites displayed various morphological abnormalities. They failed to establish normal apical polarity, and the developing protrusion was frequently thinner, abnormally curved, or otherwise deformed compared to that of WT parasites (Figure 4E). Others developed multiple protrusions or aberrant polarity, indicative of defects in morphogenesis and elongation. Although a range of abnormal developmental forms was observed by live cell imaging at both approximately 9.5 h (corresponding to stage II to III) and 24 h post-activation (Figure 4E, Supplementary Figure 3G), most *sfa1*-KO parasites (∼60-70%) failed to progress beyond stage II-III of ookinete development. In addition, although a few *sfa1*-KO ookinetes had apparent normal morphology, they were significantly less motile than WT control parasites (*P* < 0.001, Figure 4F).

Next, we investigated parasite development within the mosquito. Mosquitoes were infected with either WT-GFP or *sfa1*-KO parasites, and oocyst development was monitored over 21 days. WT-GFP parasites established a robust infection within the mosquito midgut, whereas there was a pronounced defect in *sfa1*-KO parasite development (Figure 4G). Counting oocysts on days 7, 14, and 21 revealed that there were significantly fewer *sfa1*-KO than WT-GFP oocysts (*P* < 0.001, Figure 4H). Morphological analysis revealed that *sfa1*-KO oocysts were significantly smaller than WT-GFP oocysts throughout development (*P* < 0.001, Figure 4I), suggesting a persistent defect in oocyst growth and maturation. By day 14, WT-GFP oocysts had undergone extensive sporulation, whereas *sfa1*-KO oocysts frequently failed to form well-defined sporoblasts or displayed limited and abnormal sporozoite differentiation (Figure 4G). By day 21, sporulation was largely complete in WT oocysts, while markedly abnormal sporulation patterns were displayed by *sfa1*-KO oocysts, consistent with a sustained defect in oocyst maturation and sporozoite development.

Sporozoite production in mosquito midguts was analysed at day 14. Compared to WT sporozoites, *sfa1*-KO sporozoites had severe morphological abnormalities, including loss of normal apical polarity, pronounced tapering at both ends, and excessive elongation (Figure 4J). On enumeration we have found *sfa1*-KO-infected mosquitoes contained significantly fewer midgut sporozoites than mosquitoes infected with WT controls (*P* < 0.001, Figure 4K). The ability of sporozoites to colonise the salivary glands was measured by isolating and counting sporozoites from dissected salivary glands. No sporozoites were detected in the salivary glands of *sfa1*-KO-infected mosquitoes, whereas WT-infected controls yielded readily detectable sporozoites (*P* = 0.0011), indicating major defects in transmission-stage development and/or migration through the mosquito (Figure 4L, Supplementary figure 3H).

To determine whether *sfa1*-KO parasites could be transmitted to the mammalian host, we conducted mosquito bite-back experiments with naïve mice. With WT-infected mosquitoes a patent blood-stage parasitaemia was detected 3-4 days after the mosquito feed, but mice exposed to *sfa1*-KO-infected mosquitoes developed no patent infection (Figure 4M). Finally, to determine the parental origin of the transmission defect, we performed genetic crosses between *sfa1*-KO parasites with either the male-defective (*hap2*-KO) and female-defective (*dozi*-KO) lines (*24, 25*). Examination of mosquito midguts at day 14 post-infection revealed complete restoration of oocyst sporulation following crossing with *hap2*-KO parasites (Figure 4N, O). In contrast, crossing with *dozi*-KO parasites produced only small, non-sporulating oocysts (Figure 4N, O). Consistent with these observations, mosquito bite-back experiments resulted in patent blood-stage infections (after 4 days) following the *hap2*-KO cross, but not the *dozi*-KO cross (Figure 4M). These results indicate that the transmission defect is inherited through the female line and that *sfa1*-KO male gametes retain the ability to contribute to successful transmission.

Together, these findings demonstrate that SFA1 is required for normal parasite differentiation post-fertilization in the mosquito midgut and progression through the subsequent mosquito-stages. In the absence of SFA1, defects during early ookinete morphogenesis and polarity severely compromised ookinete motility There is reduced oocyst growth and defects in sporulation, leading to disruption of sporozoite morphology and salivary gland colonisation. Overall, parasite transmission through the mosquito is ablated.

### SFA1 is required for correct establishment of ookinete polarity, pellicle formation and cytoskeletal organisation

To investigate the ultrastructural basis of the developmental defects observed in *sfa1* KO ookinetes, WT and *sfa1*-KO parasites were fixed at 9.5 and 24 h post-activation and analysed by SIM and U-ExM. SIM imaging of ookinetes stained with P28-specific antibody revealed several morphologically defective forms of *sfa1-*KO parasite at both time points (Figure 5A). Expanded samples were stained with anti-α-tubulin to display microtubule organisation and with NHS ester to reveal overall cellular architecture at high resolution. At 9.5 h post-activation, stage III WT ookinetes displayed ordered subpellicular microtubule arrays associated with the developing apical region, and MTOC differentiation associated with nuclear spindle formation. In contrast, *sfa1*-KO ookinetes had pronounced defects in apical organisation. The developing protrusion was frequently abnormally pointed, elongated, or curved, and the subpellicular microtubules appeared as a disorganised mesh rather than forming an ordered array. Distinct apical polar and basal ring structures were not readily identifiable, and in some cells more than one site of apparent apical polarity was present, with microtubules extending from two separate regions. The rounded region also frequently contained densely packed and disorganised microtubules (Figure 5B). At 24 h post-activation, WT ookinetes were mature, with a well-organised subpellicular microtubule architecture associated with the apical polar region, four clearly differentiated MTOCs, and micronemes located toward the apical end of the cell. In contrast, *sfa1-*KO parasites appeared to have been arrested at approximately stages III to IV, with an abnormally elongated or deformed protrusion. They frequently displayed dual polarity and failed to establish the ordered microtubule structures characteristic of mature WT ookinetes (Figure 5B).

**Figure 5.**
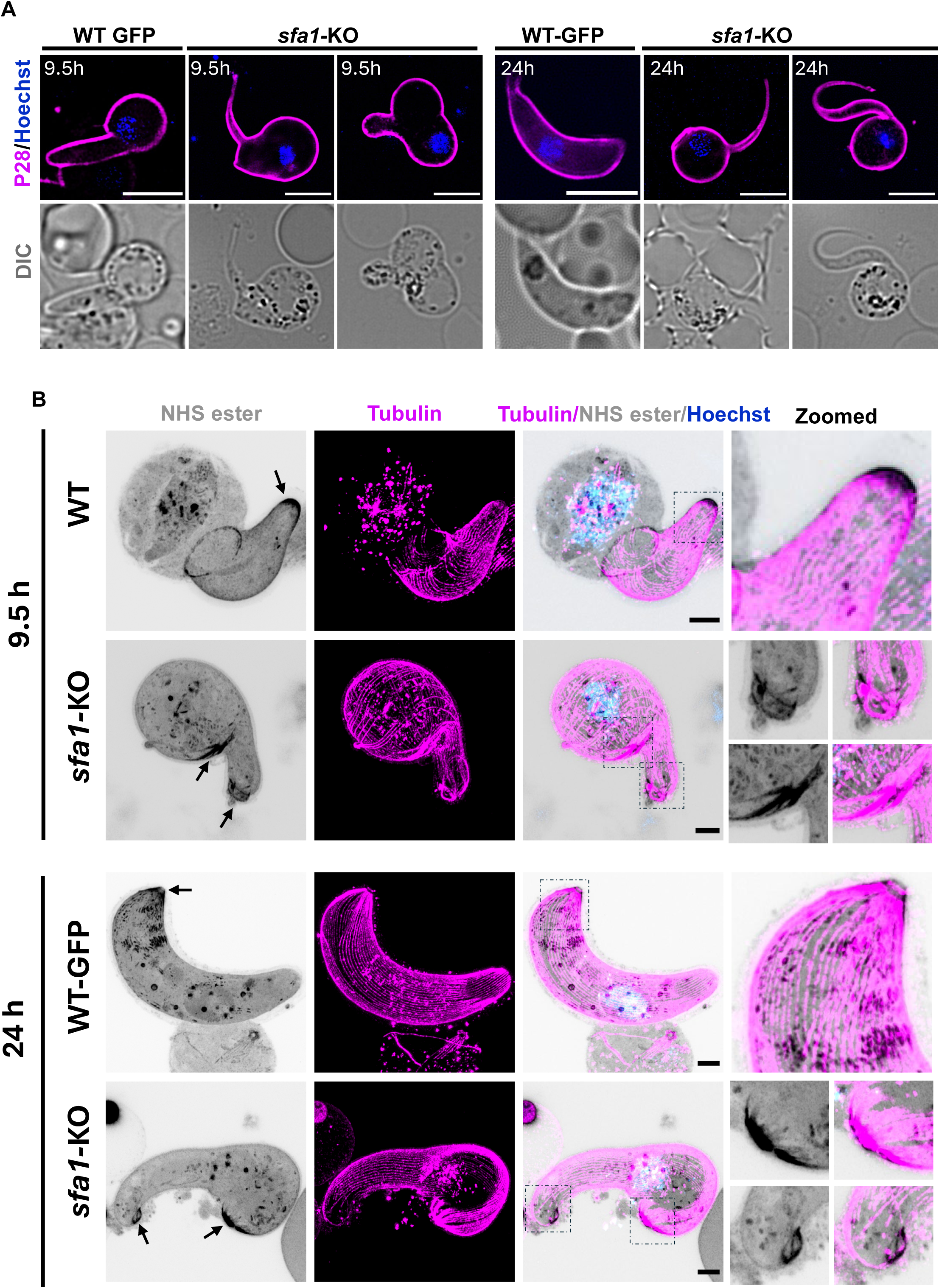
Ultrastructure of WT GFP and *sfa1-*KO ookinetes: **(A)** SIM imaging of WT-GFP and *sfa1**-***KO ookinetes fixed at 9.5 and 24 hpa. The parasites were stained with P28 antibody (magenta) and Hoechst (blue). n>20, three independent experiments. Scale bar 5 µm**. (B)** U-ExM imaging of WT-GFP and *sfa1-*KO ookinetes fixed at 9.5 and 24 hpa. The parasites were co-stained with anti-tubulin antibody (magenta), NHS ester dye (grey) and Hoechst (blue). The top panel represents the ookinetes fixed at 9.5 hpa and the bottom panel represents parasites fixed at 24 hpa. The image on the right shows the zoomed portion highlighted in black squares. n>10, three independent experiments. Arrow indicates the apical polar regions. Scale 5 µm.

In addition to expansion microscopy, similar samples were examined by transmission electron microscopy. The WT showed similar morphology to that reported previously (*23, 26*) with a single conical protrusion associated with the formation of the anterior of the developing ookinete (Figure 6A). At this stage (9.5 hours) certain nuclei showed the spindle characteristic of meiosis (Figure 6A, insert). The protrusion progressed to form the crescentic shaped mature ookinete (Figure 6B) with the apical complex consisting of a polar ring, IMC and underlying microtubules (Figure 6B, insert). In cross section there appear to be 53 even spaced microtubules running longitudinally along the length of the ookinete. In contrast, in *sfa1-*KO lines many retort forms retained the spherical shape of the zygote, while a few exhibited one or more cytoplasmic projections (Figure 6C). Unlike the WT, *sfa1-*KO parasites showed multiple areas of IMC and subpellicular microtubule (SPMT) formation (Figure 6D) and a few showed some small protrusions of the IMC (Figure 6D and E). In a few cases these formed “finger-like” projections, which in cross section showed the plasmalemma and IMC but disorganised microtubules (Figure 6C, insert and Figure 6F). Within the cytoplasm, organelles characteristic of the ookinete (micronemes and crystalline bodies) developed in the cytoplasm of the retort (Figure 6C, D).

**Figure 6.**
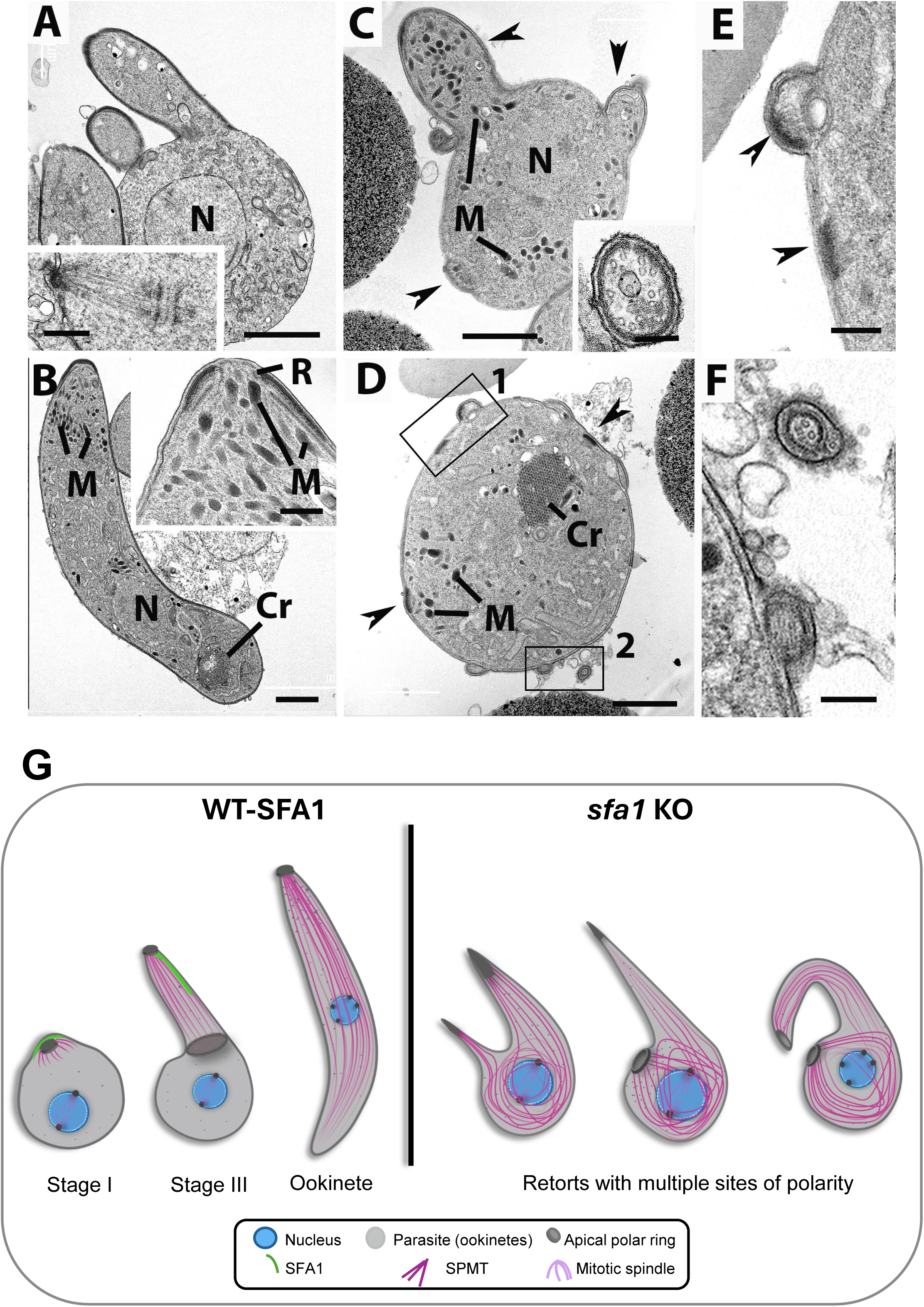
Electron microscopy analysis of WT-GFP and *sfa1-*KO ookinetes. **(A)** Early-stage WT-GFP retort form (9.5 hours post activation) showing the protrusion of the developing ookinete and the nucleus (N). Bar is 1μm **Insert.** Part of a nuclear spindle with microtubules radiating from the pole. Bar is 100nm. **(B)** Mature crescentic shaped WT-GFP ookinete showing the apical micronemes (M) plus nucleus (N) and crystalline body Cr) more posteriorly located. Bar is 1μm. **Insert.** Enlargement of the apical complex showing the anterior polar ring (R), rings form the IMC and underlying microtubules. Thin ducts run from the micronemes (M) to the apical plasmalemma. **(C)** *sfa1*-KO retort form at 9.5 hrs showing three areas with ookinete-like protrusions (arrowheads) while the cytoplasm contains a nucleus (N) and a number of micronemes (M), which are also present in the protrusions enclosed by an IMC. Bar is 1 um. **Insert.** Cross section through a finger like projection showing the plasmalemma, IMC and disorganised microtubules. Bar is 100 nm. **(D)** A spherical *sfa1*-KO retort (9.5 hrs) showing a number of electron densities associated with the plasmalemma (arrowheads and enclosed areas) while the cytoplasm contains a few micronemes (M) and a crystalline structure (Cr). Bar is 1 um. **(E)** Detail of area 1 in (D) showing electron densities associated with plasmalemma (arrowheads), one protruding from the surface. Bar is 100nm. **(F)** Detail of area 2 in (D) showing one surface protrusion and a cross section of a protrusion with an IMC and central microtubules. Bar is 100 nm. **(G)** Cartoon representation of the WT and *sfa1-*KO ookinetes, showing the stage I, stage III and mature ookinete morphology of WT parasites, and various morphological forms observed in the *sfa1*-KO parasite, including complete loss of apical polarity and having multiple polarities.

This abnormal morphology in the absence of SFA1 indicates that this protein is required for the normal subcellular spatial relationships between the apical polar region, the subpellicular microtubule array, and the elongating ookinete body (Figure 6G). The cytoskeletal organisation is disturbed in the absence of SFA1.

## Discussion

We have investigated the spatiotemporal dynamics and function of the rootlet striated fibre assemblin protein SFA1 by during replication and differentiation of the major invasive zoite stages of *P. berghei*, including merozoites, ookinetes, and sporozoites. While a recent study (*13*) described SFA1 localisation during schizogony in *P. falciparum* and sporogony in *P. berghei*, our work further complements and advances those studies with particular emphasis on the sexual zygote-to-ookinete differentiation and development, which was not previously studied. We provide a comprehensive analysis of SFA1 across multiple developmental stages during the life cycle using complementary cell biology, live cell imaging, ultrastructural microscopy and proteomics techniques. The ookinete stage is unique since it encompasses post-zygotic meiosis and the establishment of apical-basal polarity during early zygote development and where extensive cytoskeletal morphogenesis takes place, processes that are critical for successful transmission through the mosquito. Moreover, the ookinete is morphologically and biochemically distinct from sexual stage gametocytes and sporozoite stages. It is also the only motile and invasive stage that is formed extracellularly, does not require invasion of a new host cell as it moves between the mosquito gut cells while traversing to the basal surface and also develops extracellularly into the oocyst (*10, 14*).

Our live-cell imaging data confirm previous observations on the location of SFA1 during asexual blood-stage parasite replication (*13*), and extends the SFA1-GFP profile during liver stages, where it is localised to replicative stages and is absent at the late stages of merosome formation. This reveals its previously unreported association with dividing nuclei during hepatic schizogony, similar to the profile observed in the erythrocytic stages in the previous study. By co-imaging SFA1-GFP and the kinetochore marker NDC80-mCherry (*26*), we examined the spatiotemporal relationship between rootlet-fibre formation and chromosome segregation during blood stage schizogony. SFA1-GFP localisation precedes kinetochore segregation, suggesting that establishment of the rootlet occurs at, or before the earliest events of daughter cell assembly, before karyokinesis and well before cytokinesis. SFA1-GFP fluorescence was absent from mature merozoites immediately prior to egress and erythrocyte re-invasion, indicating that its function is likely restricted to the initial stages of zoite morphogenesis rather than in the mature invasive form. A similar dynamic localisation was observed during sporozoite development, consistent with previous reports, and supporting a conserved role for rootlet fibres during the early establishment of polarity and zoite formation as reported recently for merozoite and sporozoite (*13*) and for ookinete in this study.

The pattern of SFA1-GFP location during sexual development, which was not studied previously had several intriguing features. SFA1-GFP was present in non-activated female gametocytes but absent from male gametocytes and gametes. This was a surprising observation since in other eukaryotes rootlet fibres are typically associated with basal bodies and flagella (*1*), where they function in basal body orientation and the assembly or stabilisation of microtubules. During *Plasmodium* male gametogenesis there is very rapid de novo assembly of basal bodies and axonemes, and the absence of SFA1 suggests distinct parasite-specific mechanisms for basal body organisation. It is possible that in this group of organisms the primary role of SFA1 has diverged from its ancestral role. Its retention in female gametocyte stages suggests a specialised role following fertilisation, in determining cell shape, cytoskeleton organisation and polarity. Additionally, it may be acting as a template for the microtubule arrangement of the pellicle in the developing ookinete. This is analogous to SAS6L, a paralogue of the ubiquitous basal body/centriole protein SAS6, which acts as a flagellar protein in trypanosomatids and is associated with the conoid in the apicomplexan *Toxoplasma*. However, in *Plasmodium,* SAS6L is only observed in the zygote to ookinete stages of development and is absent during male gametogenesis (*27, 28*).

A significant finding of this study was the identification of a SFA1-GFP structure during zygote to ookinete differentiation. Classical ultrastructure studies had not identified rootlet-like structures in ookinetes (*14-16*) and in the recent volume electron microscopy study (*29*). It is possible that the structure may be difficult to resolve by conventional electron microscopy as it resides very close to the apical pole and is attached to the pellicle and IMC, or may be highly dynamic, transient or very close to the pellicle. Rootlet has only been observed by TEM during sporogony, where the development of sporozoites takes 14 days, whereas ookinete development takes 24 hours. However, our live-cell imaging analyses identified a striking asymmetric distribution of SFA1-GFP along one side of the pellicle during the early stages of ookinete differentiation. This distribution closely resembles that of the IMC protein ISP1, a marker of early apical polarity establishment (*30, 31*). The dynamic spatiotemporal distribution of SFA1 in ookinetes mirrors the pattern observed in merozoites and sporozoites and suggests a primary function for SFA1 in the early stages of replication, morphogenesis and establishment of cell polarity. Together these observations support the idea that in apicomplexan parasites the function of rootlet fibre is to coordinate establishment of cell polarity in coordination with programmes of cell division, rather than serving as a basal-body-associated structure.

Interaction studies examined parasite proteins present during early schizogony (15-18h) and at 8-9 h post-fertilisation ookinete development stage (stages II-III), corresponding to times when SFA1-GFP was most abundant during schizont and ookinete differentiation, respectively. Consistent with the organisation of rootlet fibres in other organisms, SFA1-GFP was bound to SFA2 in both schizonts and ookinetes (*13*). However, Pearl protein, which was reported recently to interact with *P. falciparum* SFA1 is not encoded in the *P. berghei* genome (PlasmoDB, version 71) (*13*). In addition, association with DMC1, a key meiotic recombinase suggests that rootlet formation may be coordinated with meiotic progression during ookinete development, but a direct interaction is unlikely and will have be tested specifically. A conserved protein, PBANKA_122590, was detected only in the ookinete interactome at stage II-III; this is a female stage-specific protein (PlasmoDB, version 71) that may have a role in rootlet-mediated establishment of polarity in female gametocytes/zygotes. The preferential recovery of proteins associated with subpellicular microtubules and the IMC in developing ookinetes is consistent with a role for SFA1 in organising the nascent apical cytoskeleton. In particular, the identification of SPMT1 and IMC proteins is relevant given the established roles of subpellicular microtubules and apically polarised IMC proteins in ookinete morphogenesis (*30, 31*). These observations complement the localisation and knockout phenotypes presented here but do not establish direct physical interactions. SERA2 and SERA3 may represent a distinct class of SFA1-associated candidates linked to surface or secretory compartments. Their recovery warrants further investigation, especially in light of the recent identification of SERA3 at the ookinete surface (*21*).

Functional analysis of the *sfa1* gene knockout parasite revealed several phenotypes throughout the parasite life cycle. Consistent with previous studies (*13*), *sfa1*-KO parasites had reduced growth in the asexual blood stage, confirming that SFA1 contributes to efficient parasite replication. The mislocalisation of centrin (for the centriolar plaque) and tubulin for spindle dynamics detected using fixed IFA again suggests that there is a close association of SFA1 with the centriolar plaques. This is similar to the function of SFA1 in *T. gondii,* where it has an essential role in daughter cell formation (*12*). However, male gametogenesis and exflagellation were unaffected in *sfa1*-KO parasites, consistent with the absence of SFA1 from male gametes and ruling out a direct role for SFA1 in basal body assembly. These data suggest that rootlet fibre function is restricted to female-derived developmental stages and further zoite morphogenesis during ookinete development. This differs markedly from many other eukaryotes like *Paramecium* and *Tetrahymena,* where it is associated with flagella or ciliary movement, and basal bodies (*3, 32, 33*) or *Chlamydomonas* (*34*) with abnormal basal body number in the SFA mutant (*35*).

The most severe defects resulting from *sfa1* deletion were manifested during ookinete differentiation and further parasite development in the mosquito. Profound abnormalities in cell morphology, particularly at stage II-III, were observed, including multiple polarity foci, abnormal pellicle and severe cytoskeletal disorganisation. U-ExM revealed major defects in subpellicular microtubule organisation and apical architecture, including an aberrant cell shape and the formation of several apical-like structures in individual cells as observed both by expansion and TEM analyses. These data strongly suggest that SFA1 is required to define and maintain a single apical pole during ookinete development. In the absence of SFA1, apical polarity cues are lost, preventing correct assembly of the pellicle and cytoskeleton. The function of SFA1 appears to be more restricted to cytoskeletal organisation and establishment of apical polarity at this stage. This may reflect the unique nature of ookinete development, which differs from other stages in that it does not arise through multiplication within a coenocyte such as a schizont or oocyst, nor does it involve the formation of a parasitophorous vacuole following invasion of the gut wall (*16, 17*). Although a small proportion of *sfa1*-KO parasites developed into morphologically ‘normal’ ookinetes, most arrested as abnormal retort forms. The defects in establishment of polarity and organisation of the cytoskeleton during ookinete development had profound consequences for parasite transmission. Oocyst formation was severely diminished, most likely because the abnormal ookinetes were unable to migrate efficiently and invade the mosquito midgut epithelium. The few oocysts that did form generated few sporozoites, which themselves had cytoskeletal abnormalities. The inability of mutant parasites to produce viable sporozoites resulted in complete ablation of parasite transmission to the mammalian host, complementing similar work reported for sporogony (*13*).

Taken together, our findings establish rootlet SFA1 as a key coordinator of parasite replication, establishment of apical polarity and cytoskeletal organisation of the pellicle during *Plasmodium* development. The findings suggest that the role of SFA1 in organising apical polarity is conserved across multiple *Plasmodium* zoite stages. Rootlet appears to have evolved a new specialised role in coordinating chromosome segregation, establishing cell polarity, and assembling the pellicle in *Plasmodium*, rather than functioning as a basal-body-associated structure, as described in various other eukaryotes (*1*). This new function is particularly important during ookinete development, since accurate establishment of a single apical polarity and cytoskeleton organisation with subpellicular microtubules and the IMC is essential for proper ookinete differentiation and successful mosquito transmission. Future studies to investigate the functional relationships between SFA1, IMC assembly proteins, and the newly identified ookinete-specific interaction partners will provide further insight into the evolution, structure and function of cytoskeleton-organising centres in apicomplexan parasites.

### Material & Methods

### Ethics statement

All animal procedures used in this work were reviewed and approved by the United Kingdom Home Office. Experiments were carefully conducted under the UK Home Office Project Licenses 30/3248, PDD2D5182 and PP3589958, in compliance with the *Animals (Scientific Procedures) Act 1986*. Female CD1 outbred mice (6-8 weeks old; Charles River Laboratories) were used for all experiments. Mice were subjected to 12 h light and 12 h dark (7pm till 7am) light cycle, at room temperature between 20 and 24°C, and humidity between 40 and 60%.

### Routine *P. berghei* culture and parasite lines

To begin parasite infection, mice were injected intraperitoneally (i.p.) with approximately 1 x 10^6^ blood-stage parasites. Parasite growth was monitored daily by examining Giemsa-stained thin blood smears. For erythrocytic stages, *P. berghei*-infected blood was routinely cultured in RPMI-1640 (Gibco) medium containing 1% w/v sodium bicarbonate, and 20 % v/v heat-inactivated foetal bovine serum (FBS) and a 1× penicillin/streptomycin (Sigma) antibiotic cocktail. The culture was gassed with a 5% CO₂/O₂/N₂ mixture for 2 min and maintained for 8-24 hours at 37 °C on a shaker set at 90 rpm to obtain different erythrocytic stages. For ookinete stages, blood containing gametocytes was cultured for 24 h in ookinete medium containing RPMI-1640, 100 μM xanthurenic acid, 1% w/v sodium bicarbonate, and 20 % v/v heat inactivated FBS (*36*). The NDC80-mCherry line was previously generated in our lab (*26*).

### Generation of transgenic parasites and genotype analyses

To generate C-terminally GFP-tagged transgenic lines for SFA1 (PBANKA_1227000), a region downstream of the ATG start codon of the gene was amplified, ligated into the p277 vector, and transfected as previously described (*37*). The p277 vector includes a human DHFR cassette, providing resistance to pyrimethamine. Supplementary Figure 1A shows schematic representations of the endogenous gene locus, the vector constructs, and the recombined gene locus. To confirm integration of the GFP-tagging constructs for parasites expressing C-terminal GFP-tagged protein, diagnostic PCR was performed with primer 1 (Int T260) and primer 3 (ol492) (Supplementary Figure 1B). Supplementary table 1 summarises the primers used to generate the mutant parasite lines. The pBS-DHFR plasmid was used to create the gene-deletion targeting vector for *sfa1*. This plasmid contains a *T. gondii* dhfr/ts expression cassette, which provides resistance to pyrimethamine, and is flanked by polylinker sites as described previously (*38*). PCR primers N1421 and N1422 were used to amplify a 1094 bp fragment of the 5′ sequence upstream of *sfa1* from genomic DNA, which was then inserted into the *Apa*I and *Hin*dIII restriction sites upstream of the dhfr/ts cassette in the pBS-DHFR plasmid. A 776 bp fragment from the 3′ flanking region of *sfa1* was generated with primers N1423 and N1424 and inserted downstream of the dhfr/ts cassette using the *Eco*RI and *Xba*I restriction sites. To confirm deletion of the *sfa1* gene, a diagnostic PCR using primer 1 (Int142_5) and primer 2 (ol248) was conducted (Supplementary Figure 1B and Supplementary Table 1). The *P. berghei* ANKA line 2.34 (for GFP-tagging), and ANKA line 507cl1 expressing GFP (for gene deletion) were transfected by electroporation as described previously (*39*).

### Purification of schizonts and gametocytes

To purify schizonts, blood cells collected from infected mice (day 4 post-infection) were cultured in schizont medium for 15-18h at 37°C (rotating at 90 rpm). Schizonts were purified the following day on a 60% v/v NycoDenz gradient in 1x phosphate-buffered saline (PBS) (NycoDenz stock solution: 27.6% w/v NycoDenz in 5 mM Tris-HCl (pH 7.20), 3 mM KCl, 0.3 mM EDTA).

Gametocytes were purified as described previously with slight modifications (*40*). Briefly, parasites were injected into mice pre-treated with phenylhydrazine and gametocytes enriched by sulfadiazine treatment 2 days post-infection. Blood was collected 4 days post-infection, and gametocyte-infected cells were purified using a 48% NycoDenz gradient (prepared with NycoDenz stock solution as above). Gametocytes were harvested from the gradient interface and washed thoroughly in 1x PBS.

### Live-cell imaging

To investigate the expression and location of SFA1-GFP during asexual development, parasites cultured in schizont medium were imaged at the various erythrocytic stages of development. Purified gametocytes were assessed for GFP expression and location at different time points (0 and 1 to 15 min) post-activation in ookinete medium. Zygote and ookinete stages were labelled using the Cy3-conjugated mouse monoclonal antibody 13.1 (red), which targets the P28 surface protein (*19*) and examined over a 24-hr period. Oocysts and sporozoites were imaged in infected mosquito guts. All images were captured with a 63x oil immersion objective on a Zeiss AxioImager M2 microscope equipped with an AxioCam ICc1 digital camera.

### Western blot analysis

To detect SFA1-GFP in schizont lysates, purified schizonts were lysed for 30 min in a buffer containing 10 mM Tris-HCl (pH 7.5), 150 mM NaCl, 0.5 mM EDTA, and 1% NP-40. Samples were sonicated for 1 min at 40% amplitude (1 sec on, 1 sec off). The samples were centrifuged at 15,000*g* at 4 °C for 20 min, and the supernatant was diluted with buffer containing 10mM Tris/Cl pH 7.5, 150mM NaCl, 0.5 mM EDTA and incubated with GFP-Trap Agarose beads (Chromotek) for at least 1 h with end- to-end rotation. Beads were pelleted and washed thoroughly in wash buffer (10mM Tris/Cl pH 7.5, 150mM NaCl, 0.05% Nonidet P40). Bead samples were boiled at 95 °C for 5 min in Laemmli buffer and centrifuged at 13000 × g for 5 min. Bead supernatant samples were separated on a 4-12% SDS-polyacrylamide gel, and the resolved proteins were transferred onto a nitrocellulose membrane (Amersham Biosciences). Immunoblotting was performed using the Western Breeze Chemiluminescence Anti-Rabbit kit (Invitrogen) with an anti-GFP polyclonal antibody (Invitrogen) diluted at 1:1,250, following the manufacturer’s instructions.

### Parasite phenotype analyses

Blood samples containing approximately 1,000 *sfa1*-KO parasites were injected intraperitoneally (i.p.) into mice. The development of asexual-stage parasites and gametocyte production were monitored by microscopy of Giemsa-stained thin blood smears. Four to five days post-infection, gametocytes were harvested, and exflagellation and ookinete conversion were examined on a Zeiss AxioImager M2 microscope. To analyse mosquito infection and parasite transmission, 30 to 50 *Anopheles stephensi* SD 500 mosquitoes were allowed to feed for 20 min on anaesthetised, infected mice with at least 15% asexual parasitaemia and a comparable gametocyte level. To examine midgut infection, approximately 15 guts were dissected from mosquitoes on days 7 and 14 post-feeding, and oocysts were counted by microscopy using a 63× oil immersion objective. On day 21 post-feeding, another 20 mosquitoes were dissected, and their guts and salivary glands were crushed separately in a loosely fitting homogeniser to release sporozoites, which were then quantified using a haemocytometer or used for imaging. Mosquito bite-back experiments with naïve mice were conducted 21 d post-feeding, and blood smears were examined after 3 to 4 days. For genetic complementation, we used either male (*hap2-*KO) or female (*dozi-*KO) defective parasites using a method described previously(*36*) Briefly, mice were infected with combinations of the different parasite lines mentioned above and subsequently fed to mosquitoes. These were dissected 14 dpi and the number of live or dead oocysts determined as mentioned above.

### Immunoprecipitation and mass spectrometry

Immunoprecipitation was performed for purified schizonts (15-18h post culture) to obtain dividing cells, gametocytes (non-activated) and ookinete pellets (stage III) from SFA1-GFP parasites. Wild-type GFP (WT-GFP) schizonts, non-activated gametocytes and stage III ookinetes were used as controls. The respective parasites were cross-linked with 1% formaldehyde for 10 min, followed by a 5 min incubation in 0.125 M glycine solution. Samples were washed three times in PBS. Cell lysates were prepared, and immunoprecipitation was conducted using a GFP-Trap_A Kit (Chromotek) according to the manufacturer’s instructions. Briefly, purified cells were lysed for 30 min in a buffer containing 10 mM Tris-HCl (pH 7.5), 150 mM NaCl, 0.5 mM EDTA, and 1% NP-40. Samples were sonicated for 1 min at 40% amplitude (1 sec on, 1 sec off). The samples were centrifuged at 15,000 × g at 4°C for 20 min, and the supernatant diluted with buffer containing 10mM Tris/Cl pH 7.5, 150mM NaCl, 0.5 mM EDTA. To capture SFA1-GFP and bound interactors, samples were incubated with GFP-Trap Agarose beads (Chromotek) for at least 2 h with end-to-end rotation at 4°C. Beads were pelleted and washed thoroughly in wash buffer (10mM Tris/Cl pH 7.5, 150mM NaCl, 0.05% Nonidet P40) to remove unbound proteins. On-bead tryptic-digestion peptides were analysed by liquid chromatography-tandem mass spectrometry. Mascot (http://www.matrixscience.com/) and MaxQuant (https://www.maxquant.org/) search engines were used for mass spectrometry data analysis. Experiments were performed in triplicate as described in (*41*). Peptides and proteins with a minimum threshold of 95% were used for further proteomic analysis. The PlasmoDB database (version 71) was used for protein annotation, and manual curation was used to classify SFA1-GFP interactions as biologically relevant.

For the analyses shown in Figure. 3, precursor intensity for each protein was divided by the sum of protein precursor intensities in the corresponding MS run and multiplied by 10. Values were log₂-transformed before comparison. Proteins detected in fewer than two retained SFA1-GFP preparations per stage were excluded from the corresponding volcano analysis. Following quality-control exclusions, analyses included four schizont, three ookinete and two gametocyte SFA1-GFP runs, with four schizont and three gametocyte WT-GFP control runs; the three gametocyte WT-GFP controls were also used for the ookinete comparison due to low amount of protein recovery in the ookinete controls. The latter are shown in Supplementary Table 2 for comparison. Protein-level differences between groups were evaluated using Welch’s t-test (two-sided), and results were visualised as ΔPI (mean log₂ intensity in SFA1-GFP minus mean log₂ intensity in the selected GFP control) against −log₁₀(P) in a volcano plot. Missing values were inputted by random sampling of values around ΔPI=10. P-values were corrected for multiple testing using Benjamini-Hochberg. FDR was set to 5%; 10% for gametocytes.

### Fixed Immunofluorescence assay

Purified *sfa1*-KO schizonts were smeared onto poly-L-lysine-coated slides and fixed with ice-cold 1:1 v/v acetone/methanol for 5 min. Immunocytochemistry used anti-GFP, anti-α-tubulin mAb (Sigma-T9026; used at 1:1000) or anti-centrin (Sigma-mouse monoclonal, clone:20H5, 04-1624; used at 1:500) as the primary antibodies, and the secondary antibody was Alexa 488 (Invitrogen-A1101) or 568-conjugated IgG for the respective primary antibodies (Invitrogen-A11004) (used at 1:1000). Slides were mounted in Vectashield with DAPI (Vector Labs) for fluorescence microscopy with a Zeiss AxioImager M2 microscope fitted with an AxioCam ICc1 digital camera.

### Liver stage cultures

Human liver hepatocellular carcinoma (HepG2) cells (European Cell Culture Collection) were grown in DMEM medium, supplemented with 10% heat-inactivated FBS, 0.2% NaHCO_3_, 1% sodium pyruvate, 1% penicillin/streptomycin and 1% L-glutamine in a humidified incubator at 37°C with 5% CO_2_. For infection, 1 × 10^5^ HepG2 cells were seeded in each well of a 48-well culture plate. The day after seeding, sporozoites were purified following mechanical disruption of salivary glands harvested from female *A. stephensi* mosquitoes infected with SFA1-GFP parasites and added to the infection medium. The culture was maintained for 72 hr and used for immunofluorescent assays.

### Ultrastructure expansion microscopy (U-ExM)

Purified schizonts, activated gametocytes or ookinetes were fixed in 4% paraformaldehyde (PFA, Sigma) diluted in microtubule (MT)-stabilising buffer (MTSB) (*41-43*). Fixed cells were allowed to settle on a 12 mm diameter poly-D-lysine (A3890401, Gibco) coated coverslip for 2 h at room temperature. Coverslips were incubated overnight in 1.4% formaldehyde (FA)/2% acrylamide (AA) at 4°C. Gelation and expansion were performed as described previously (*44*). Immunolabelling was performed using primary antibodies against mouse α-tubulin (1:1000; Sigma, T9026), rabbit GFP (1:250; Thermo Fisher), or mouse centrin (1:500; Thermo Fisher). Secondary antibodies were anti-rabbit Alexa Fluor 488 (Invitrogen) and anti-mouse Alexa Fluor 568 (Invitrogen), each used at 1:1000 dilution. For bulk proteome labelling, the gels were stained with Atto 594 NHS or Atto 665 (Merck) diluted to 10 µg/ml in PBS, for 90 min at room temperature on a shaker. Images were acquired on a Zeiss Elyra PS.1-LSM780 system and CD7-LSM900, and subsequent image analysis was performed using Zeiss ZEN 2012 (Black edition) and Fiji/ImageJ (version 1.54p).

### Structured illumination microscopy

Small volumes (3 µl) of schizont, gametocyte or ookinete parasite suspensions were co-stained with 13.1-conjugated antibody against the ookinete surface protein P28. Cells were placed on a microscope slide and covered with a long (50 × 22 mm) coverslip to obtain a very thin monolayer and immobilise the cells. Cells were scanned with an inverted microscope using a Zeiss Plan-Apochromat 63x/1.4 oil immersion objective on a Zeiss Elyra PS.1 microscope, utilising the structured illumination microscopy (SIM) technique. The SIM mode settings included lasers: 405nm: 20%, 488nm: 8-16%; 561 nm: 15%-40%, 642 nm: 20% exposure times: 300 or 500 ms (Hoechst), 300 ms (GFP); 80-300 ms (NDC80-mCherry, P28, Tub), 300 ms (Centrin) five grid rotations; five phases. The bandpass filters BP 420-480 + LP 750, BP 495-550+LP 750, BP 570-620 and LP 655 were used for the blue, green, red and far-red channels, respectively. Where multiple focal planes (Z-stacks) were recorded, processing and channel alignment were done as described previously (*36*).

### Electron microscopy

Ookinetes or gametocytes activated for 8 min and 15 min were fixed in 4% glutaraldehyde in 0.1 M phosphate buffer and processed for electron microscopy as described previously (*45*). Briefly, samples were post-fixed in osmium tetroxide, treated en bloc with uranyl acetate, dehydrated, and embedded in Spurr’s epoxy resin. Thin sections were stained with uranyl acetate and lead citrate prior to examination in a JEOL JEM-1400 electron microscope (JEOL, United Kingdom). The experiments were performed at least three times to capture every stage of the parasite, and 50 to 55 cells were examined for phenotypic analysis.

### RNA isolation and quantitative real-time PCR (qRT-PCR) analyses

RNA was extracted from purified schizonts or gametocytes using an RNA purification kit (Stratagene). RNA-to-cDNA kit (Applied Biosystems) was used to synthesise complementary DNA (cDNA). SYBR Green Fast Master Mix kit (Applied Biosystems) was used to quantify gene expression from 80 ng of total RNA. All primers were designed using Primer3 (Primer-BLAST, NCBI). The analysis was conducted on an Applied Biosystems 7500 Fast machine with the following cycling conditions: 95°C for 20 s, followed by 40 cycles of 95°C for 3 s and 60°C for 30 s. Three technical replicates and three biological replicates were performed for both wild-type GFP and *sfa1*-KO lines. The genes *hsp70* (PBANKA_081890) and *arginyl-tRNA synthetase* (PBANKA_143420) were used as endogenous control reference genes. The primers used for qPCR are listed in Supplementary Table 1.

### Statistical analysis

Data are presented as mean ± SD or SEM, as indicated in the figure legends. Group comparisons used unpaired t-tests, a two-tailed Mann–Whitney U test for ookinete motility, or two-way ANOVA, as appropriate. Proteomic data were normalised, log₂-transformed and analysed using Welch’s t-test. P < 0.05 was considered significant; False Discovery Rate was set between 5-10% for the proteomics (Benjamini-Hochberg). ns indicates not significant. Sample sizes and numbers of independent experiments and exact p values are provided in the figure legends. All analyses and graph generation were performed using GraphPad Prism 10.

## Data availability

The mass spectrometry proteomics data have been deposited in the ProteomeXchange Consortium via the PRIDE partner repository, and data are available with identifier. Source data and the protein-level tables supporting Figure 3 are available in Supplementary Table 2 (MS-MS data).

---

**Supplementary Figure 1:**
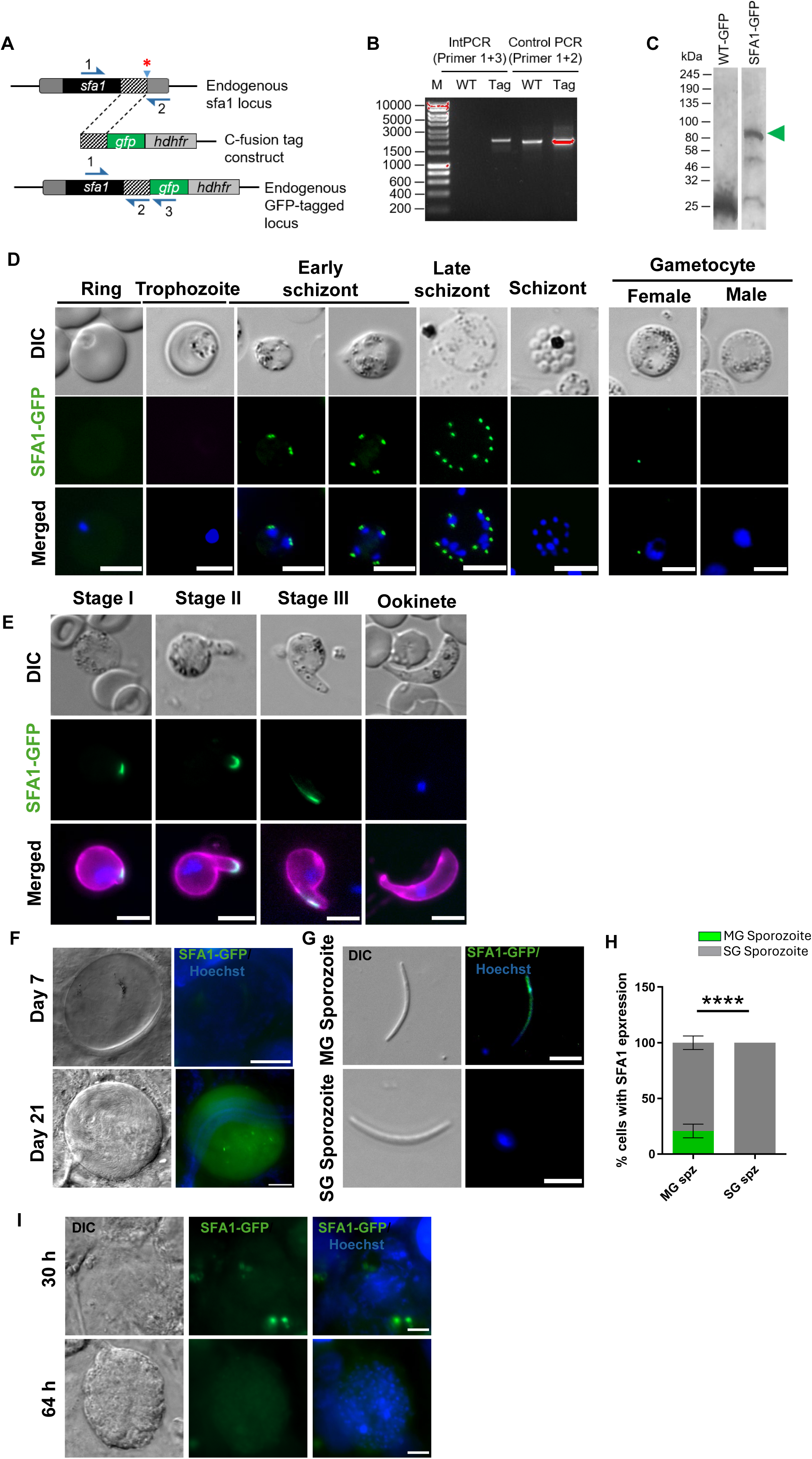
Generation, validation, and subcellular localisation of SFA1-GFP in *Plasmodium berghei.* **(A)** Schematic representation of SFA1-GFP tagging construct design showing the endogenous *sfa1* locus, the fusion tag and the recombined GFP-tagged locus. *shows the stop codon. **(B)** PCR verification of the successful insertion of SFA1-GFP and WT-GFP as a negative control (lanes 1 and 2), as well as a control PCR for both SFA1-GFP and WT-GFP parasite DNA (lanes 3 and 4). **(C)** Western blot analysis showing GFP expression in WT-GFP (25kDa) and SFA1-GFP (∼ 95 kDa) from purified schizonts. **(D)** Live-cell images of SFA1-GFP blood-stage parasites showing the location in schizonts and in male and female gametocytes. Scale bar 5 µm **(E)** Live-cell images of SFA1-GFP across the developmental stages of ookinetes. Scale bar 5 µm. **(F)** SFA1-GFP location in oocysts at day 7 and day 21. Scale bar 10 µm. **(G)** SFA1-GFP localisation in 21-day midgut and salivary gland sporozoites. Scale bar 5 µm. **(H)** Quantification of the proportion of sporozoites showing expression of SFA1-GFP in midguts MG and salivary gland SG sporozoites (n = 50 parasites; graph indicates mean ± SD). Two-way ANOVA,****p<0.0001, three independent experiment **(I)** SFA1-GFP (green) expression in early (30 h) and late (64h) exo-erythrocytic forms (liver stages). Hoechst (blue). Scale bar 5 µm.

**Supplementary Figure 2:**
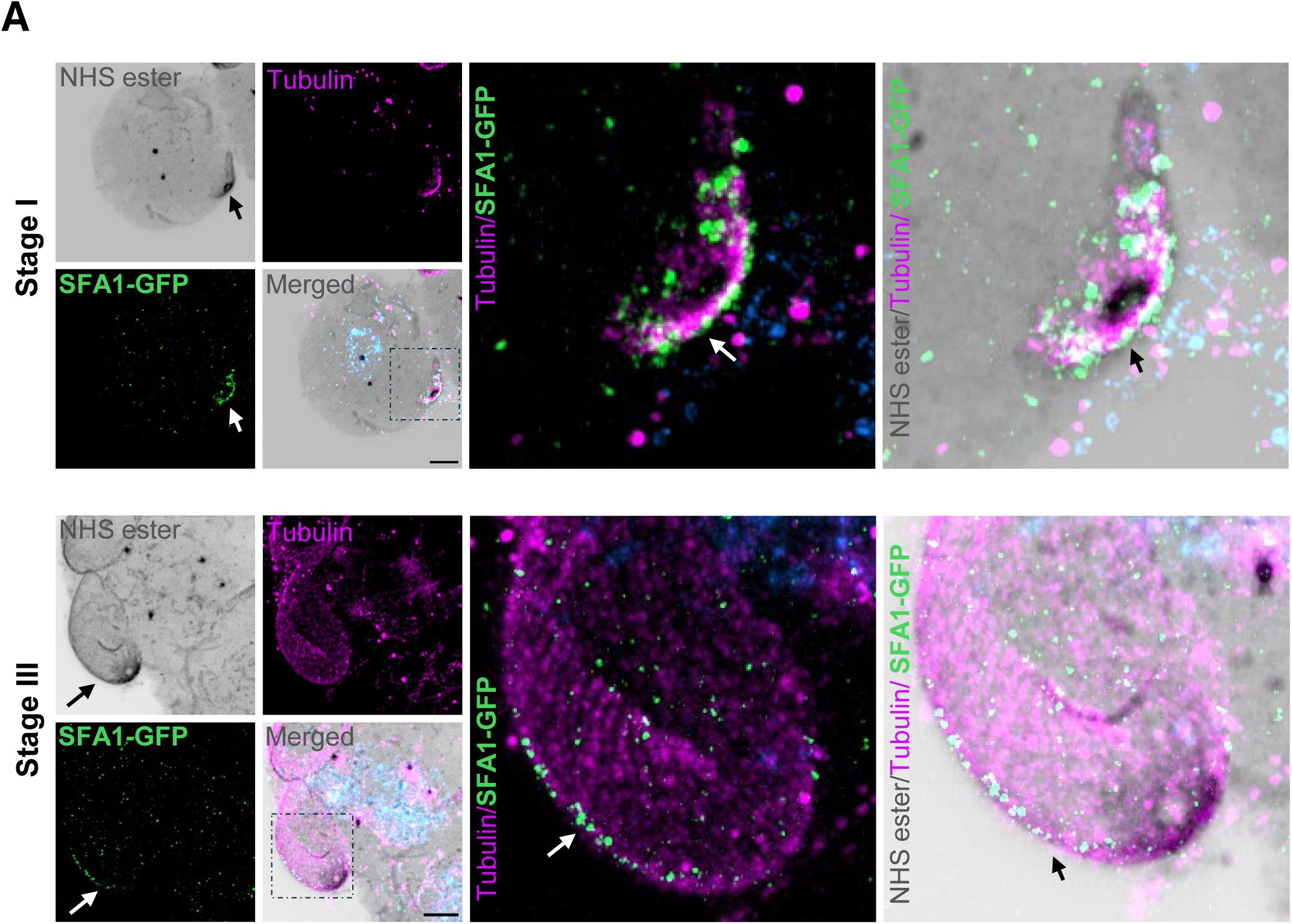
Subcellular location of SFA1 in zygotes: **(A)** U-ExM of SFA1-GFP ookinete stages showing SFA1-GFP detection in stage I (top panels) and stage III (bottom panels). Ookinetes were co-stained with anti-GFP (green), anti-tubulin (magenta) to visualise microtubules, and NHS ester (red) to broadly label cellular ultrastructure. Scale bars 5 µm. The inset shows the magnified view of the boxed region. Arrow indicates the apical polar regions and localisation of SFA1-GFP (green).

**Supplementary Figure 3.**
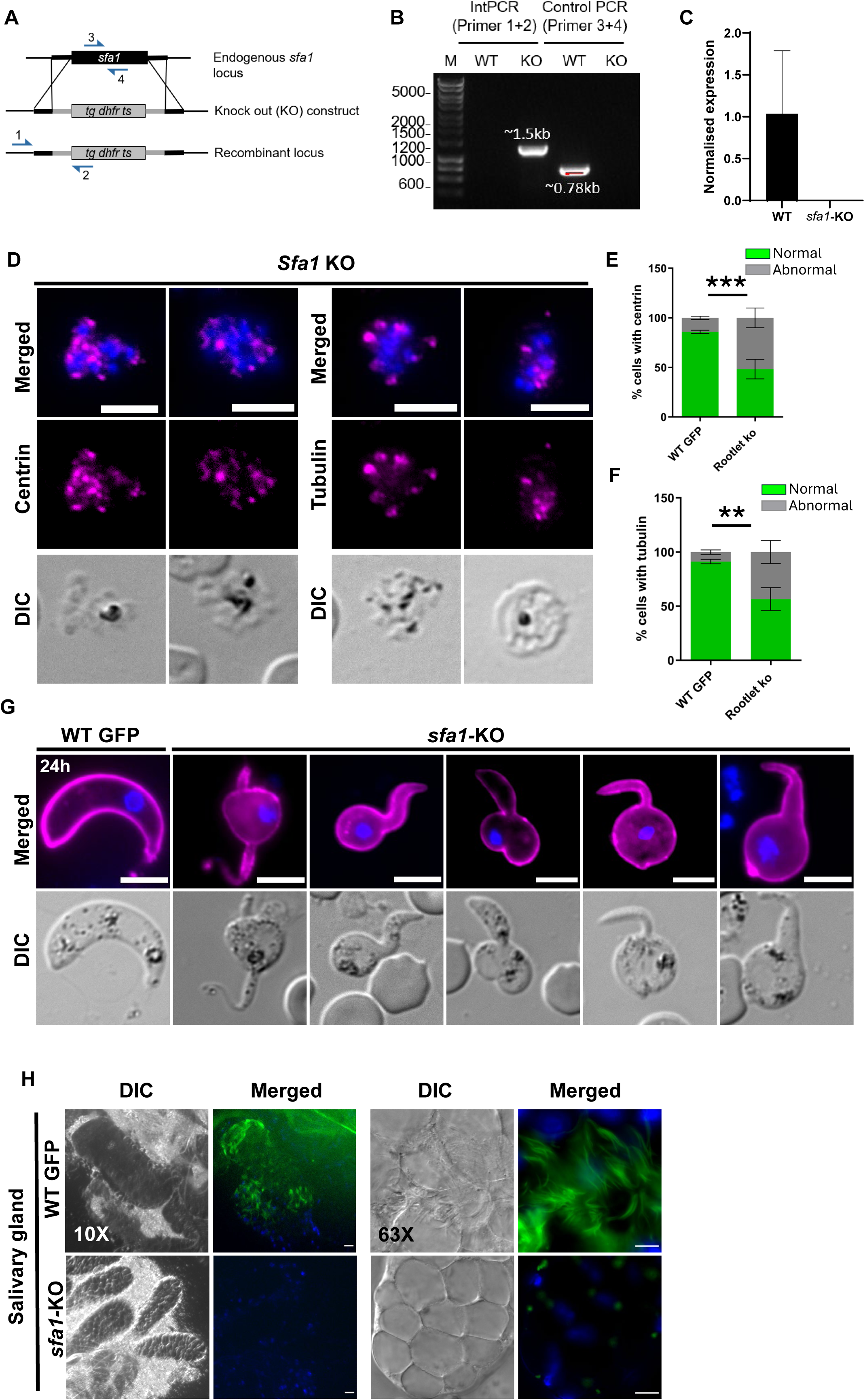
Generation of *sfa1* knockout line and its phenotypic analysis. **(A)** Schematic representation of the endogenous *sfa1* locus, the targeting gene deletion construct, and the recombinant locus after double homologous recombination. **(B)** Integration PCR showing correct integration with expected band sizes and deletion of the *sfa1* gene from the knockout (KO) parasite line. **(C)** *sfa1* gene expression in WT-GFP and *sfa1*-KO parasite lines. Mean ± SEM. n=3 technical replicates and 3 biological replicates. **(D)** Immunofluorescence imaging of *sfa1* KO blood stage schizonts co-stained with anti-centrin (magenta) and anti-tubulin (magenta) antibody, and Hoechst (blue). **(E)** Quantification of the proportion of asexual blood stage parasite showing deformed organization of centrin (n > 150 parasites; graph indicates mean ± SD, Two-way ANOVA),***p=0.0007, three independent experiment. **(F)** Quantification of the proportion of asexual blood stage parasite showing deformed organization of tubulin (n > 150 parasites; graph indicates mean ± SD, Two-way ANOVA), **p<0.0019, three independent experiment. **(G)** Live-cell imaging showing the ookinete morphology of WT-GFP and *sfa1-*KO lines. In magenta is a cy3-conjugated antibody, 13.1, which binds to the P28 protein on the surface of ookinetes. Panels: DIC (differential interference contrast); merged: Hoechst (blue, DNA) and P28 (magenta). Scale bar = 5 μm. **(H)** Live cell images of mosquito salivary glands containing the WT-GFP and *sfa1**-***KO parasite lines (Scale bar 10 µm).

**Supplementary Figure 4.**
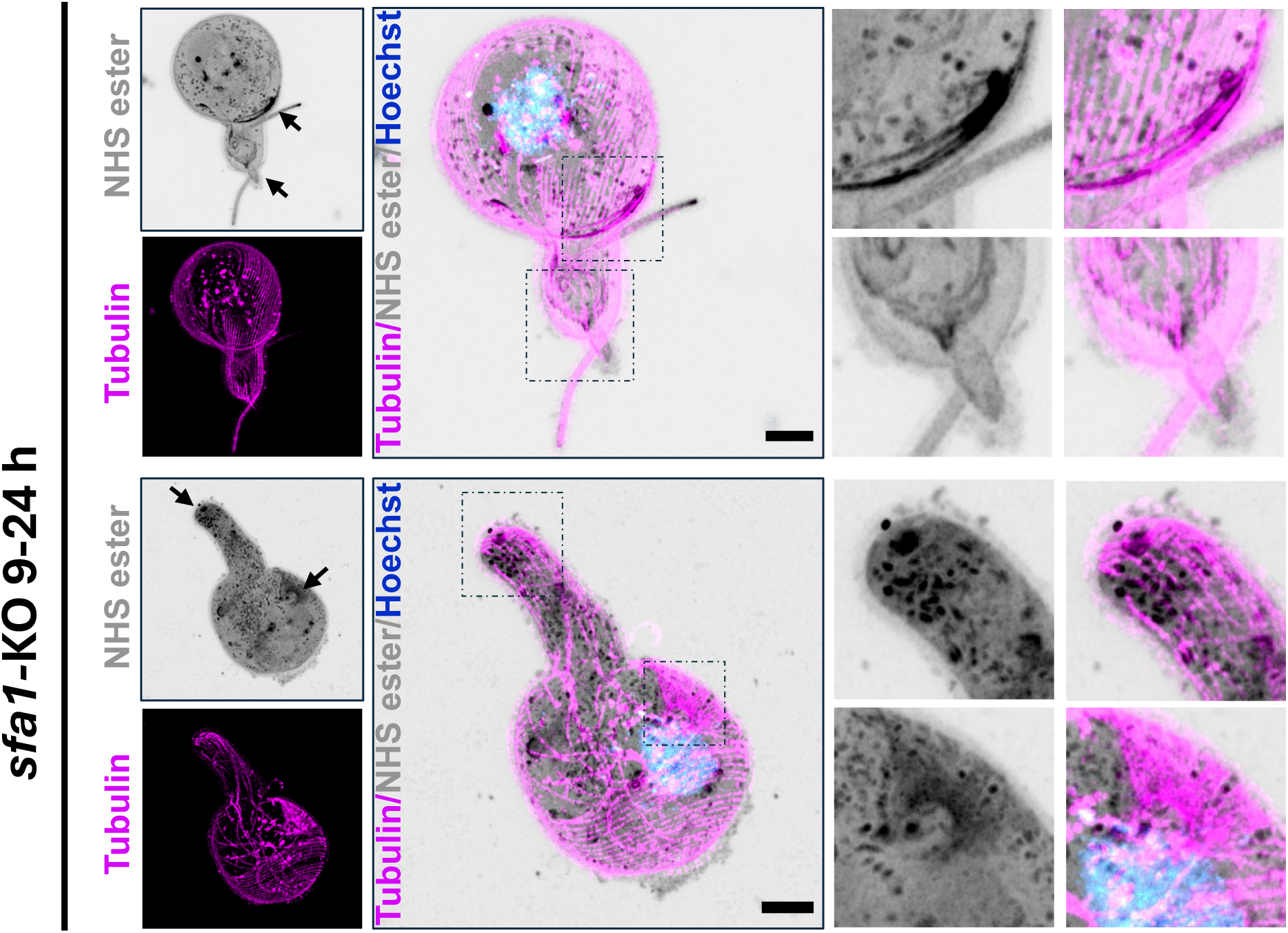
Ultra structural imaging of *sfa1-*KO ookinetes. UExM images showing *sfa1**-***KO ookinetes co-stained with anti-tubulin (magenta) to visualise microtubules, and NHS ester (red) to label protein-dense cellular structures. Scale bars 5 µm. The inset shows the magnified view of the boxed region. Arrow indicates the apical polar regions.

**Supplementary table 1:**
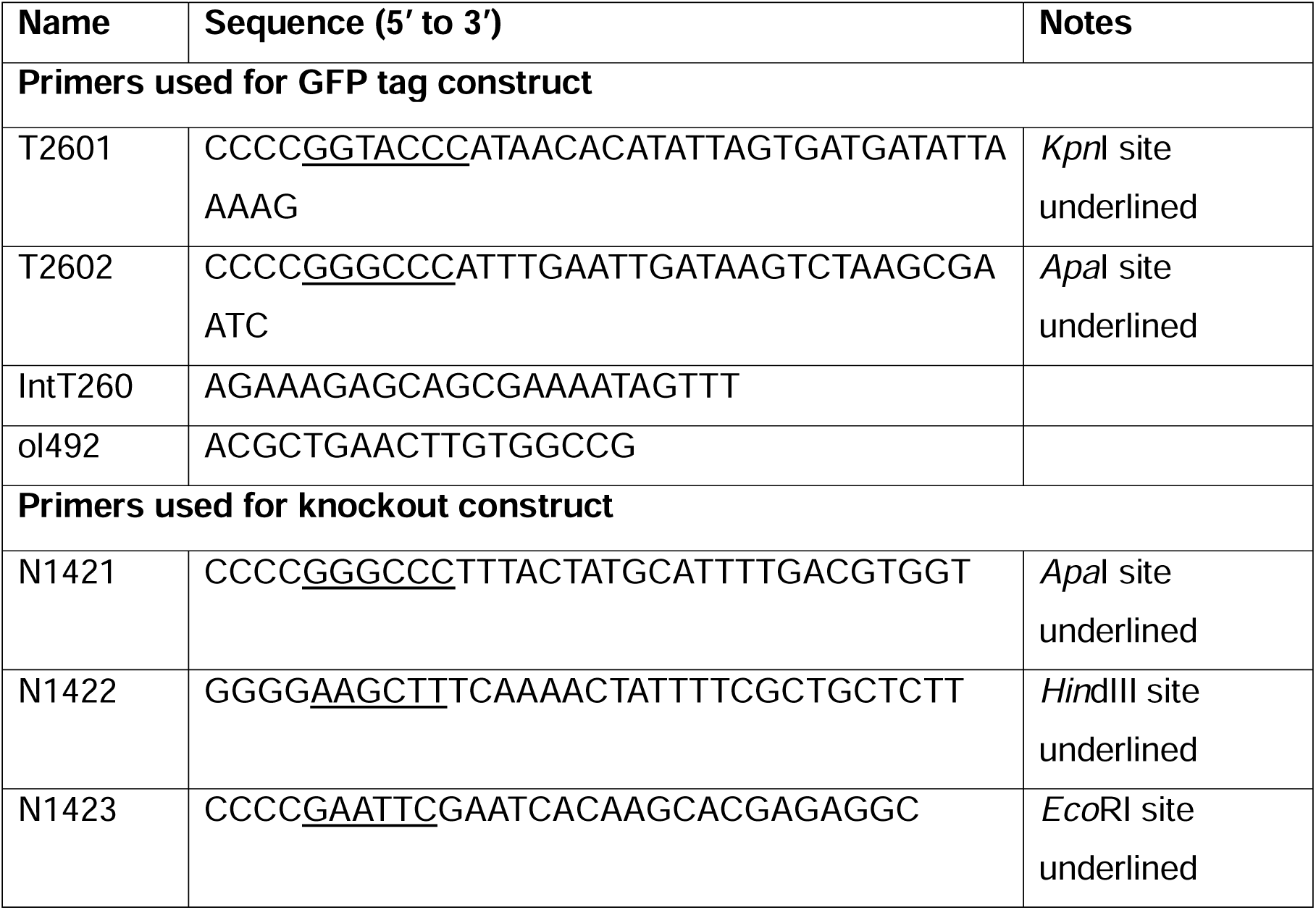
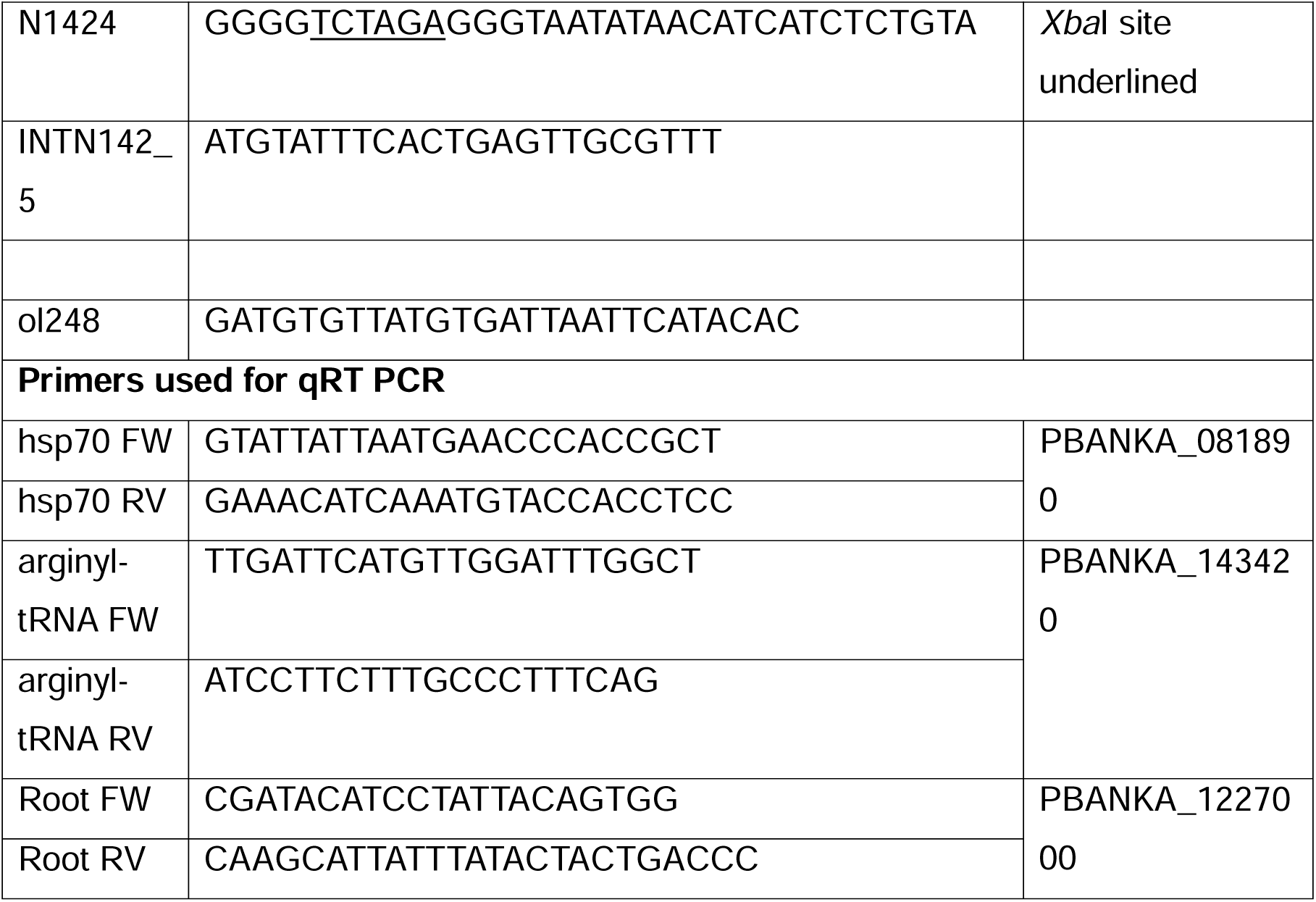
Oligonucleotides used in this study.

Supplementary Table 2: MS-MS overview.

## Author Contributions

RT conceived the project. RT, DB, AM, PI, DSG, MZ, performed the live cell imaging, genotypic and phenotype analysis; DJPF, SV performed TEM analysis; RT, AM and RM performed the SIM microscopy; AM and PI performed U-ExM observation; RT, AM, ARB and ECT performed proteomic analysis; AM, PU performed the liver stage experiments: DJPF, ECT, AAH, AM, PI, SV, DSG, RT performed formal analysis; RT acquired funding and supervised the project. AM, PI and RT wrote the first manuscript draft; AAH edited the first draft. All authors reviewed and edited the manuscript.

## Funding and Acknowledgement

RT is supported by an ERC advance grant funded by UKRI Frontier Science (EP/X024776/1), MRC UK (MR/K011782/1), and BBSRC (BB/L013827/1, BB/X014681/1). AM, PI, PU, MZ, were supported as research fellows and DB as senior technician (EP/X024776/1). DSG is supported by the BBSRC (BB/X014681/1). AAH is supported by the Francis Crick Institute (FC001097), which receives core funding from the Cancer Research UK (FC001097), the UK Medical Research Council (FC001097), and the Wellcome Trust (FC001097). ECT was supported by a personal fellowship from the Nederlandse Organisatie voor Wetenschappelijk Onderzoek (NWO), the Netherlands (grant no. VI. Veni.202.223). We thank the School of Life Sciences Imaging (SLIM) for providing access to confocal and SIM microscopy. We thank Cleidiane Zampronio at Warwick University for mass spectrometry methods and Bio Support Unit, University of Nottingham for maintenance of mice used in this study. For Open Access, the authors have applied a CC BY public copyright licence to any Author Accepted Manuscript version arising from this submission.

